# Protein abundance and folding rather than the redox state of Kelch13 determine the artemisinin susceptibility of *Plasmodium falciparum*

**DOI:** 10.1101/2021.07.02.450839

**Authors:** Robin Schumann, Eileen Bischoff, Severina Klaus, Sophie Möhring, Julia Flock, Sandro Keller, Kim Remans, Markus Ganter, Marcel Deponte

## Abstract

Decreased susceptibilities of *Plasmodium falciparum* towards the endoperoxide antimalarial artemisinin are linked to mutations of residue C580 of Kelch13, which is the homologue of the redox sensor Keap1 in vertebrates. Here, we addressed whether mutations alter the artemisinin susceptibility by modifying the redox properties of Kelch13 or by compromising its native fold or abundance. Using selection-linked integration and the *glmS* ribozyme, efficient down-regulation of Kelch13 resulted in ring-stage survival rates around 40%. While the loss of a potential disulfide bond between residues C580 and C532 had no effect on the artemisinin suceptibility, the thiol group of C473 could not be replaced. We also established a protocol for the production of recombinant Kelch13. In contrast to cysteine-to-serine replacements, common field mutations resulted in misfolded and insoluble protein. In summary, not the redox properties but impaired folding of Kelch13, resulting in a decreased Kelch13 abundance, is the central parameter for mutant selection.

## Introduction

Artemisinin and its derivates are central to the treatment of malaria and are recommended as first-line drugs for artemisinin-based combination therapies by the World Health Organization^1^. In 2009, Dondorp et al. reported a delayed clearance of the malaria parasite *Plasmodium falciparum* in patients following artesunate treatment^2^. The delayed parasite clearance was found to be associated with reduced drug susceptibility of the ring stage^3–6^ as well as mutations in *PFKELCH13*, which encodes a kelch domain-containing protein on chromosome 13^7–9^. PfKelch13 belongs to the top 5% of most conserved proteins in *Plasmodium* and comprises an N-terminal apicomplexan-specific region followed by a CCC domain, a BTB domain, and a six bladed kelch β-propeller domain^10^. Relevant mutations for delayed parasite clearance were predominantly found in the β-propeller domain of PfKelch13, with C580Y being the most prevalent one^7, 8^. Mutations R539T or I543T are less frequent but were reportd to lead to even higher ring-stage survival *in vitro*^9^. While decreased artemisinin susceptibilities were initially restricted to hot spots at the Thai–Cambodian border, non-related C580Y mutant strains have recently been detected in South America^11^ and on New Guinea^12^. Furthermore, a novel strain with a R561H mutation has emerged in Rwuanda^13^. Hence, PfKelch13 mutations endanger the long-term goal to eliminate malaria^14^.

N-terminally GFP-tagged wild-type and C580Y mutant PfKelch13 localize to the same punctate structures close to the digestive vacuole^15, 16^ and were found in ring-shaped cytostome-like structures at the plasma membrane^17^. Furthermore, tagged and untagged wild-type and mutant PfKelch13 variants were reported to localize to cytosolic foci, the endoplasmic reticulum, vesicular structures, and the mitochondrion^18, 19^. Mislocalization and hemoglobin uptake studies in combination with a dimerization-induced quantitative BioID PfKelch13 interactome revealed an involvment of PfKelch13 in endocytosis, suggesting that PfKelch13 mutations result in a decreased protein abundance, hemoglobin uptake, and activation of artemisinin^16, 17^. This theory has been supported so far by mislocalization and overexpression studies using protein-tagged wild-type or mutant PfKelch13^16–18^.

PfKelch13 is highly similar to Keap1^8, 10^, which is the master redox and electrophile sensor in mammals and which interacts with the transcription factor Nrf2 via its β-propeller domain^20, 21^. Keap1-bound Nrf2 becomes ubiquitinated and undergoes proteasomal degradation in the cytsosol^22, 23^. Oxidation or alkylation alters the conformation of Keap1, resulting in the liberation and translocation of Nrf2 to the nucleus^20–24^. Nuclear Nrf2 forms heterodimers and binds with its basic leucine zipper domain to the electrophile-responsive element (EpRE), resulting in a plethora of adaptive responses such as the induction of phase II detoxifying enzymes and the synthesis of glutathione^21, 25, 26^. Although *P. falciparum* blood stages are thought to adapt to numerous endogenous and environmental oxidative challenges, they lack an Nrf2 homologue^8^. Whether PfKelch13 also acts as a redox sensor (e.g., based on residue C580), and whether the endoperoxide artemisinin interferes with such a function, remained to be studied.

Here, we used selection-linked integration (SLI)^15^ in combination with *glmS* ribozyme-tagging^27^ and established a purification protocol for recombinant PfKelch13 to study the relevance of the abundance, conformational stability, and redox state of PfKelch13 for the artemisinin susceptibility in *P. falciparum*.

## Results

### Knockdown of *PFKELCH13* decreases parasite growth

To test the relevance of the PfKelch13 abundance on the growth of *P. falciparum* blood stages, we used the SLI method by Birnbaum *et al.*^15^ to generate a 3D7 strain with His_8_-tagged PfKelch13 that is under control of the *glmS* ribozyme from *Bacillus subtilis*^27^ (Fig. 1a,b). The ribozyme can be activated by the addition of glucosamine (GlcN), resulting in the degradation of the His_8_-PfKelch13-encoding mRNA. A strain with the inactive mutant *glmS M9* was generated in parallel and served as a negative control. Following successful selection with the *P. falciparum* dihydroorotate dehydrogenase inhibitor DSM1, plasmid integration and disruption of endogenous *PFKELCH13* was confirmed by analytical PCR for both strains (Fig. 1c). Furthermore, PCR analyses with a primer pair that binds to endogenous *PFKELCH13* up- and downstream of the homology region (and that does not recognize recodonized His_8_-*PFKELCH13*) confirmed the loss of the original *PFKELCH13* copy in the whole genome. The effect of GlcN was tested for asynchronous blood-stage cultures of strains His_8_-*PFKELCH13-glmS* and His_8_-*PFKELCH13-M9* (Fig. 1d). After five days, strain His_8_-*PFKELCH13-glmS* grew normal in the presence of 0.3 mM GlcN, whereas treatments with 1.0 mM and 2.0 mM GlcN resulted in growth inhibitions around 25% and more than 50%, respectively. To confirm a gene-specific knockdown effect and to exclude a growth inhibitory effect of GlcN, we treated His_8_-*PFKELCH13-M9* with the same GlcN concentrations. While 0.3 and 1.0 mM GlcN had no effect on parasite growth, a growth inhibition around 30% was detected for the negative control after five days of treatment with 2.0 mM GlcN. Complete growth arrest was observed for both strains in the presence of 5 and 10 mM GlcN, further supporting the reported toxicity of higher GlcN concentrations^28^. In summary, we established that up to 2.0 mM GlcN can be used for knockdown studies with His_8_-*PFKELCH13-glmS* and that the addition of 1.0 mM GlcN resulted in a specific growth defect because of the down-regulation of His_8_-PfKelch13.

**Figure 1.**
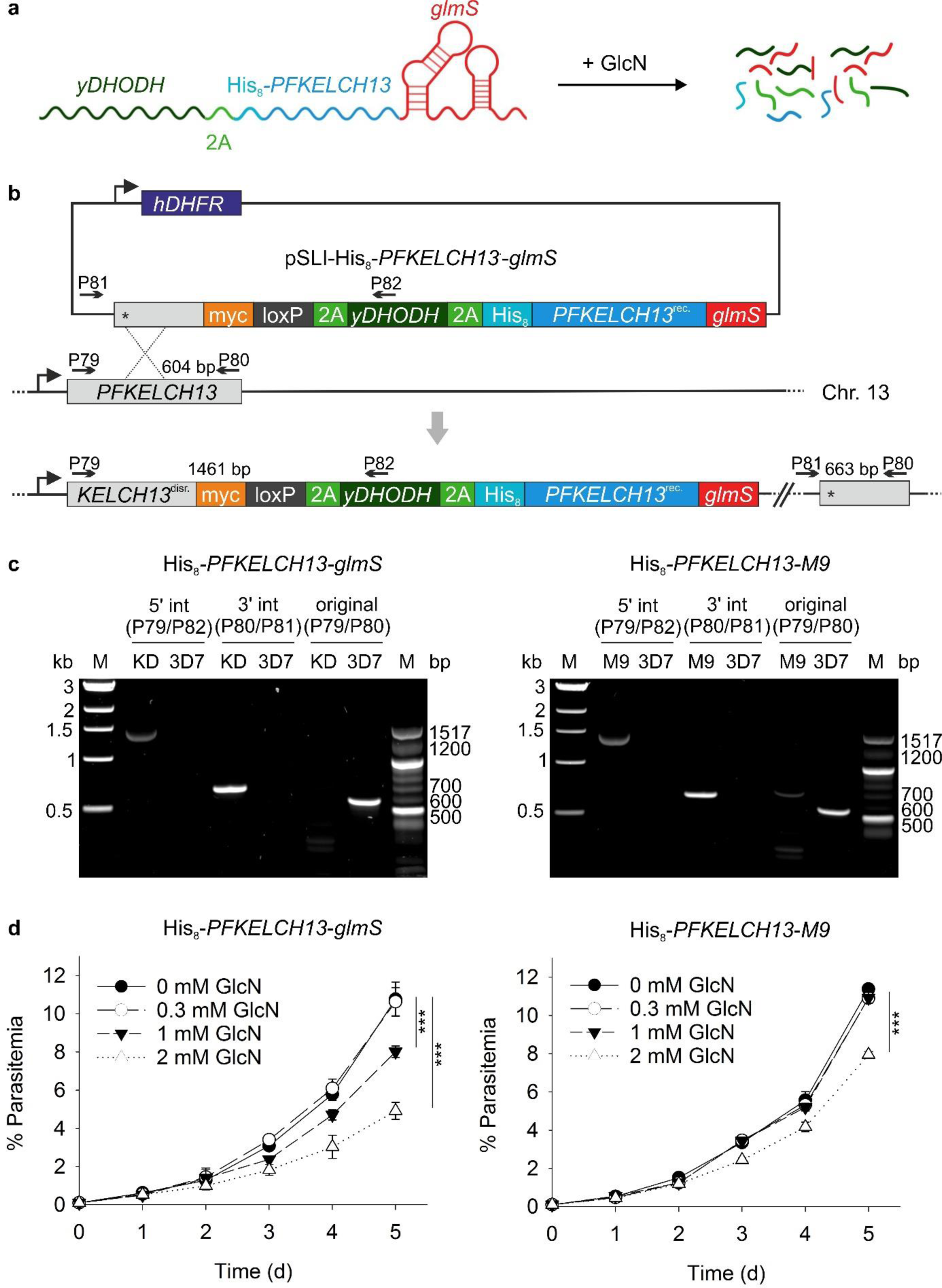
Generation and validation of a regulatable *PFKELCH13* knockdown strain. (a) Schematic overview of the knockdown strategy. The addition of glucosamine (GlcN) results in the activation of the *glmS* ribozyme and the degradation of mRNA encoding cytosolic yeast dihydroorotate dehydrogenase (*yDHODH*), the self-cleaving 2A peptide (2A) and His_8_-tagged Kelch13. (**b**) Schematic overview of the gene replacement by selection-linked integration (SLI). Plasmid-containing 3D7 parasites with human dihydrofolate reductase (*hDHFR*) are first selected using the antifolate WR99120. Disruption of the endogenous *PFKELCH13* open-reading frame by homologous recombination provides the promoter for the fusion construct between recodonized *PFKELCH13*^rec^ and *yDHODH*, the resistance marker against DSM1. Primer positions and expected product sizes for PCR analysis are highlighted. (**c**) PCR analyses using the indicated primer pairs from panel b confirmed the successful integration for knockdown strain His_8_-*PFKELCH13-glmS* (KD) and strain His_8_-*PFKELCH13-M9* (M9) with active and inactive *glmS*, respectively. Genomic DNA from parental strain 3D7 served as a control. (**d**) Growth curve analyses for ansynchronous blood-stage cultures of strain His_8_-*PFKELCH13-glmS* in the presence of different concentrations of GlcN. The parasitemia was determined from Giemsa-stained blood smears. Strain His_8_-*PFKELCH13-M9* served as a negative control to discriminate between growth defects that were caused by the knockdown of *KELCH13* or the toxicity of GlcN. All data points represent the mean ± standard deviation of three independent experiments. Statistical analyses were performed using the one-way ANOVA method in SigmaPlot13 (P ≤ 0.001: ***). Source data are provided as a Source Data file.

### Quantification of the down-regulation of His_8_-PfKelch13

The resistance marker yeast dihydroorotate dehydrogenase (yDHODH) and His_8_-PfKelch13 are encoded by the same mRNA (Fig. 1a). We therefore used the DSM1 resistance of strains His_8_-*PFKELCH13-glmS* and His_8_-*PFKELCH13-M9* as a quantitative readout for the *glmS*-dependent down-regulation of His_8_-PfKelch13 (Fig. 2a). The parental 3D7 strain served as a positive control with an EC_50_ value of 37 ± 1 nM. Strain His_8_-*PFKELCH13-glmS* showed a GlcN concentration-dependent loss of resistance towards DSM1. While 0.3 mM GlcN only slightly increased the DSM1 susceptibility, treatments with 1.0 and 2.0 mM GlcN resulted in EC_50_ values of 350 ± 30 and 116 ± 8 nM, respectively. Strain His_8_-*PFKELCH13-M9* remained fully resistant even in the presence of 2.0 mM GlcN. Thus, the loss of resistance against DSM1 confirmed the efficient *glmS*-dependent knockdown of the mRNA fusion construct between *yDHODH* and His_8_-*PFKELCH13*.

**Figure 2.**
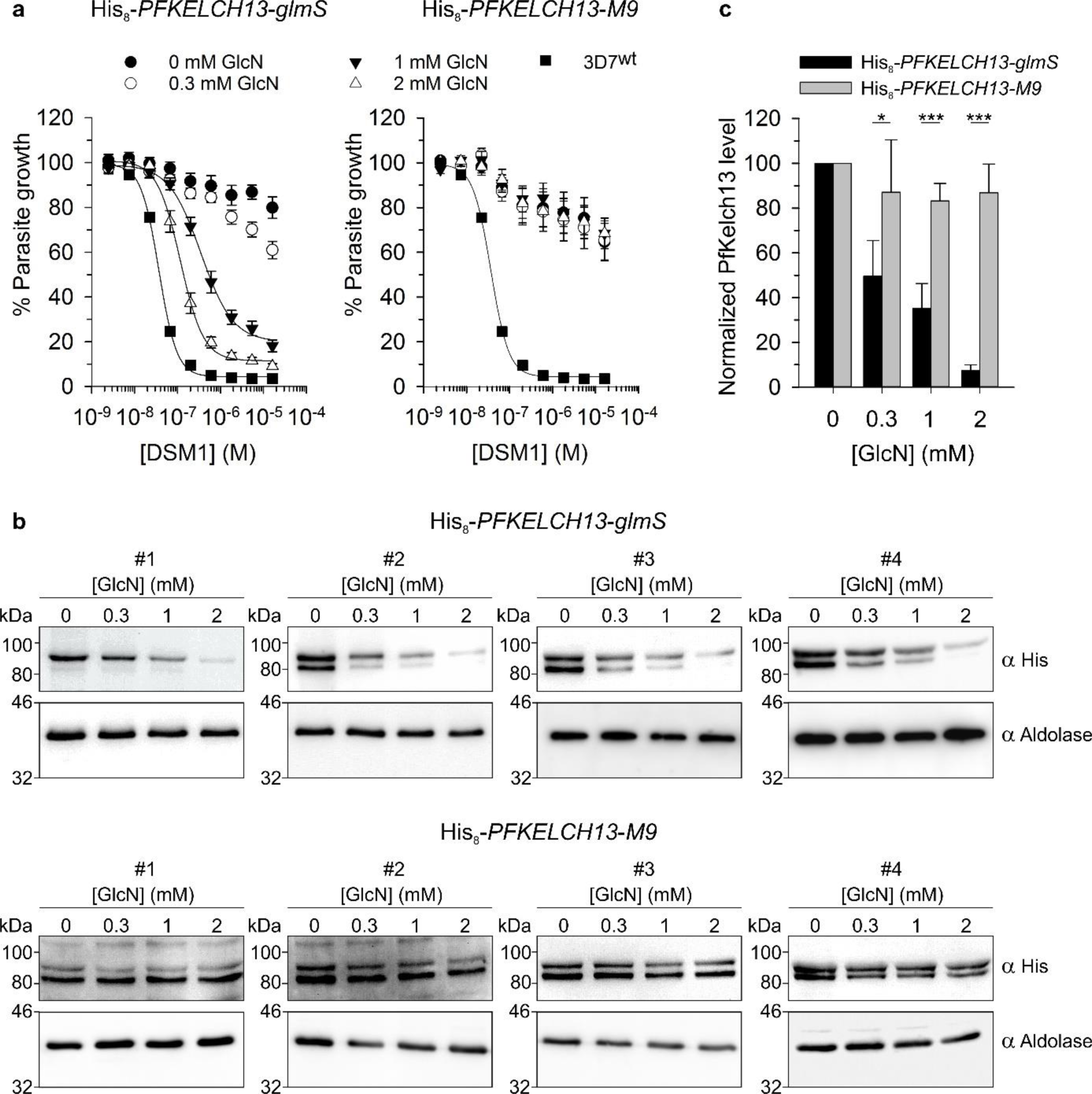
Quantification of the down-regulation of His_8_-PfKelch13. (**a**) EC_50_ values for DSM1 were determined in 96-well plates using a SYBR green assay. Ring-stage parasites from strains His_8_-*PFKELCH13-glmS* and His_8_-*PFKELCH13-M9* were incubated in 96-well plates for 140 h with or without 0.3 mM GlcN and 2.5 nM – 16.2 µM DSM1. Strain 3D7 served as a positive control. All data points represent the mean ± standard deviation of four independent experiments with triplicate measurements. (**b**) Ring-stage parasites from strains His_8_-*PFKELCH13-glmS* and His_8_-*PFKELCH13-M9* were incubated for two cycles with the indicated concentrations of GlcN before western blot analysis against the His-tag. The calculated molecular mass of His_8_-PfKelch13 is 85.2 kDa. Aldolase was used as a loading control for normalization. (**c**) Quantification of the normalized western blot signals from the four independent experiments in panel b. Statistical analyses were performed using the one-way ANOVA method in SigmaPlot13 (P < 0.05: *; P ≤ 0.001: ***). Source data are provided as a Source Data file.

Next, we performed western blot analyses against the His-tag to quantify the efficiency of the down-regulation of His_8_-PfKelch13. Four independent experiments showed a strong GlcN concentration-dependent down-regulation of His_8_-PfKelch13 (Fig. 2b). Since the antibody detected two regulated bands in parasite lysates (as outlined below), we quantified the intensity as the sum of both bands and normalized it against western blot signals for aldoalase as a housekeeping protein. In the presence of GlcN, His_8_-PfKelch13 levels decreased in a concentration-dependent manner. The addition of 1.0 mM GlcN resulted in a down-regulation of 65 ± 10%, whereas 2.0 mM GlcN resulted in a down-regulated by 93 ± 2% (Fig. 2b,c). Treatment of strain His_8_-*PFKELCH13-M9* with the same GlcN concentrations did not reveal significantly lower His_8_-PfKelch13 levels. In summary, the *glmS* system allowed us to titrate the level of His_8_-PfKelch13 with GlcN and to down-regulate the protein by more than 90%.

### Down-regulation of His_8_-PfKelch13 decreases artemisinin susceptibility

Next, we investigated whether decreased His_8_-PfKelch13 levels alter the artemisinin susceptibility comparable to PfKelch13 mutants in the field. Pretreatment of strain His_8_-*PFKELCH13-glmS* with 1.0 mM GlcN followed by a ring-stage survival assay (RSA) with artesunate without GlcN resulted in survival rates of more than 35% (Fig. 3a). Increasing the GlcN concentration during the preincubation to 2.0 mM slightly improved the survival rate, suggesting a survival rate limit around 40%, a value more than twice as high as the survival rate for the positive control NF54K13^C580Y^ at around 14%. The negative control His_8_-*PFKELCH13-M9* did not show significantly increased survival rates in the presence of GlcN. In summary, down-regulation of His_8_-PfKelch13 led to a decreased artemisinin susceptibility with a striking survival rate slightly above 40%.

**Figure 3.**
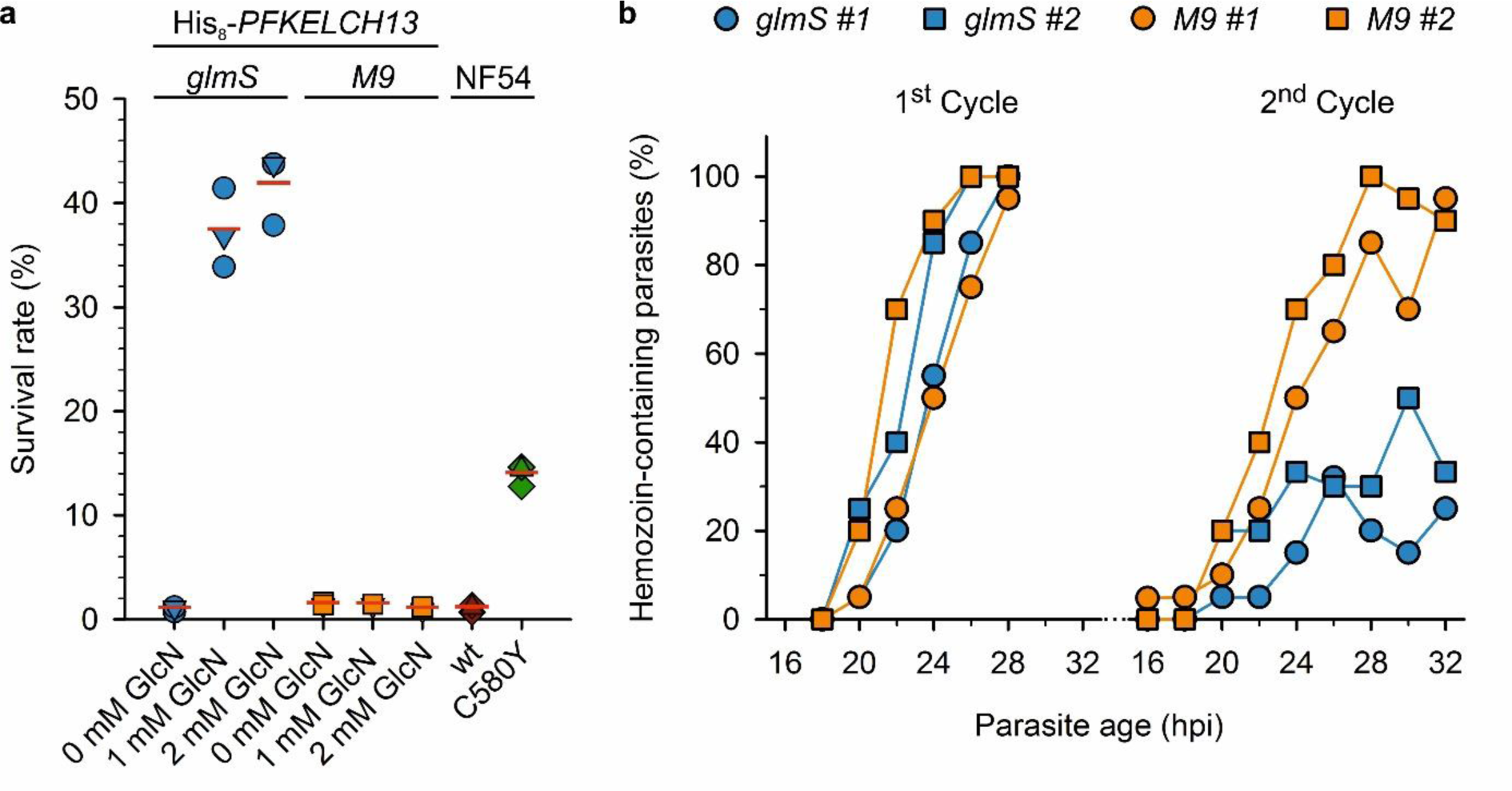
Down-regulation of His_8_-PfKelch13 increases the ring-stage survival rate and impairs hemozoin formation. (**a**) Ring-stage His_8_-*PFKELCH13-glmS* parasites (*glmS*) were pretreated for two cycles with or without the indicated GlcN concentration. RSAs were subsequently performed with 0.7 µM artesunate without GlcN. The RSAs were validated using strain NF54K13^C580Y^ as a positive control. Strains His_8_-*PFKELCH13-M9* (*M9*) and wild-type NF54 (wt) served as negative controls. Each symbol represents a data point from one of three independent experiments. (**b**) Phenotypical analysis of His_8_-*PFKELCH13-glmS* knockdown parasites. Hemozoin-containing His_8_-*PFKELCH13-glmS* (*glmS*) and His_8_-*PFKELCH13-M9* (*M9*) parasites that were treated with 1 mM GlcN were quantified during the first and second cycle 16–32 hpi. The percentage was determined by light microscopy of stained thin blood smears. Representative parasites are shown in Fig. S1. All data points represent the individual data points of two independent biological replicates. Source data are provided as a Source Data file.

### Down-regulation of His_8_-PfKelch13 impairs hemozoin formation

Parasites with a decreased artemisinin susceptibility as well as parasites with mislocalized PfKelch13 were shown to remain in a prolonged ring stage with a decreased endocytotic activity^15, 29, 30^. We therefore analyzed whether down-regulation of His_8_-PfKelch13 affects the formation of hemozoin from endocytosed hemoglobin using synchronized parasite cultures (Fig. 3b, Fig. S1). A phenotype in the presence of 1 mM GlcN was detected by light microscopy during the second intraerythrocytic cycle. While hemozoin was visible in all parasites from strains His_8_-*PFKELCH13-glmS* and His_8_-*PFKELCH13-M9* around 28 h post-invasion (hpi) during the first cycle, only around 30% of His_8_-*PFKELCH13-glmS* parasites contained visible amounts of hemozoin 28–32 hpi during the second cycle. Thus, down-regulation of His_8_-PfKelch13 phenocopied the delayed development and impaired hemozoin formation of strains with mutant PfKelch13 variants.

### Relevance of the thiol groups of C469, C473, C532, and C580

Next, we addressed whether PfKelch13 has a Keap1-like redox function and whether such a function might play a role for the artemisinin susceptibility. First, we determined the IC_50_ values of the disulfide-inducer diamide, the oxidant *tert*-butyl hydroperoxide (tBuOOH), the endoperoxide artesunate, and the reductant dithiothreitol (DTT) for strain NF54K13^C580Y^ from Ghorbal et al.^31^ (Fig. S2). Strain NF54 served as a control. The IC_50_ values were almost identical for both strains and were in good agreement with previous measurements for strain _3D7_^32, 33^.

PfKelch13 has seven cysteine residues, all of which are within the kelch β-propeller domain (Fig. 4a). Residues C532 and C580 can form a disulfide bond between two blades at the center of the β-propeller (Fig. 4b). Free access to residue C532 from the top site of the β-propeller is partially blocked by residues Y480 and N530, whereas smaller molecules might access the sulfur atom of residue C580 from the bottom site next to the BTB domain. To test the physiological relevance of the redox state of residues C532 and C580, we used SLI to generate mutant strains His_8_-*PFKELCH13^C532S^* and His_8_-*PFKELCH13^C580S^* (Fig. S3a). Strain His_8_-*PFKELCH13^wt^* with His-tagged wild-type PfKelch13 was generated as a control. PCR analysis confirmed the plasmid integration and disruption of endogenous *PFKELCH13* for all three strains (Fig. S3b). RSAs with artesunate did not reveal an altered artemisinin susceptibility for strains His_8_-*PFKELCH13^C532S^* and His_8_-*PFKELCH13^C580S^* (Fig. 4c). Strain NF54K13^C580Y^ served again as a positive control.

**Figure 4.**
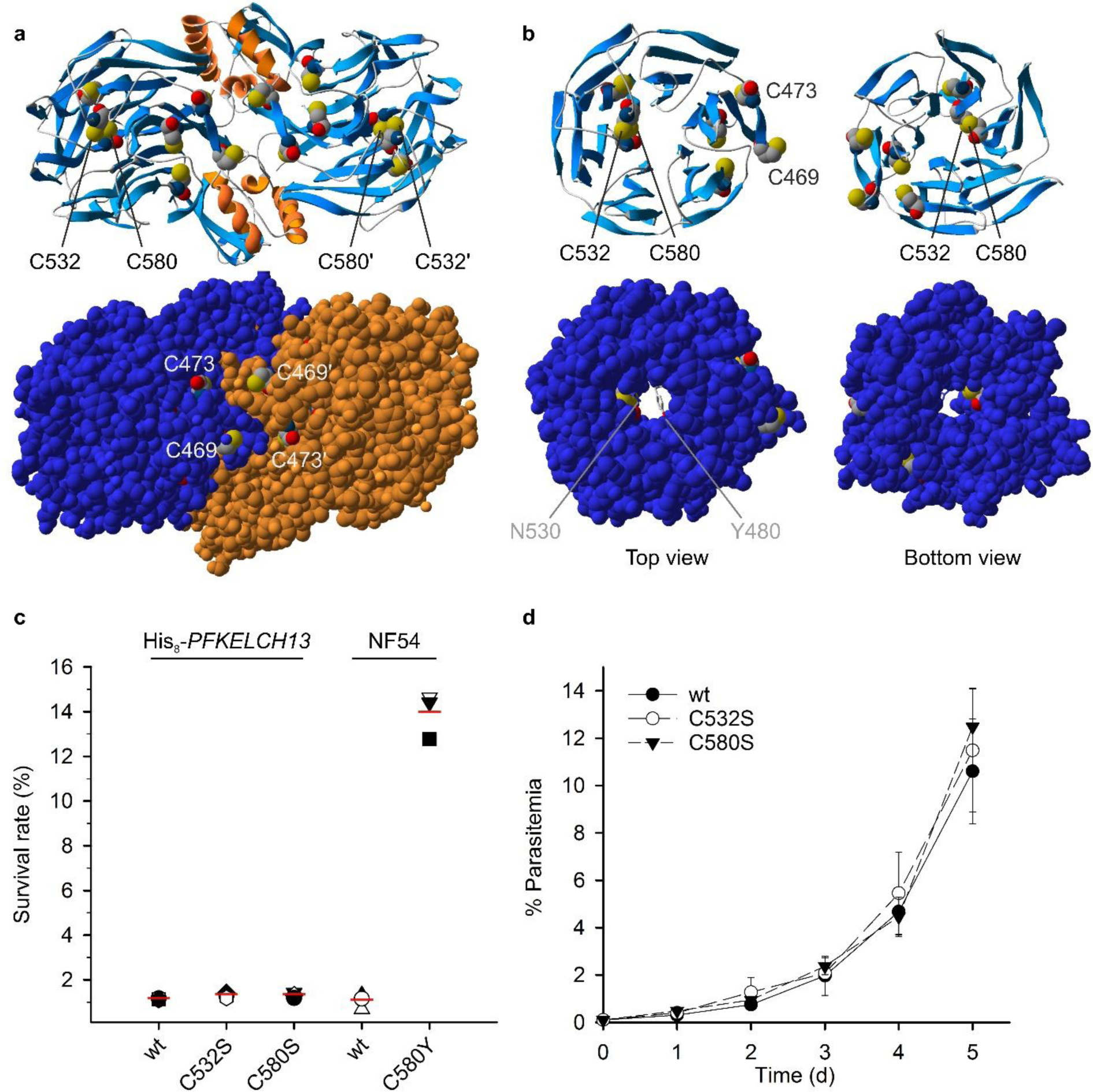
Physiological relevance of the thiol groups of C532 and C580. (**a**) Overview of the positions and accessibility of the seven cysteine residues in crystallized recombinant PfKelch^350-726^, comprising the BTB domain and the Kelch domain (PDB entry 4YY8). The two subunits of the homodimer are coloured in blue and orange in the space-filling model at the bottom. (**b**) Top and bottom view of the Kelch domain. Residues C532 and C580 can form a disulfide bond that faces the pore of the β-propeller. Residues N530 and the gatekeeper residue Y480 partially block the access of the pore from the top and are shown in stick representation. (**c**) RSAs with 0.7 µM artesunate for strains His_8_-*PFKELCH13^wt^* (wt), His_8_-*PFKELCH13^C532S^* (C532S), and His_8_-*PFKELCH13^C580S^* (C580S). Strains NF54K13^C580Y^ (C580Y) and wild-type NF54 (wt) served as controls. Each symbol represents a data point from one of three independent experiments. (**d**) Growth curve analyses for ansynchronous blood-stage cultures with His_8_-tagged PfKelch13 cysteine mutants. The parasitemia for the indicated mutant strains was determined from Giemsa-stained blood smears. Strain His_8_-*PFKELCH13^wt^* served as a control. All data points represent the mean ± standard deviation of three independent experiments. Statistical analyses were performed using the one-way ANOVA method in SigmaPlot13 (P ≤ 0.001: ***). Source data are provided as a Source Data file.

In contrast to previous reports on the generation of *PFKELCH13^C580Y^* mutants^15, 31^, we were unable to generate strain His_8_-*PFKELCH13^C580Y^*. Two out of three unsuccessful SLI experiments to generate His_8_-*PFKELCH13^C580Y^* were performed in parallel to the successful generation of other His_8_-*PFKELCH13* mutants, suggesting that the C580Y mutation in combination with the His_8_-tag and altered codon usage might be lethal. We also tried to replace the only two surface-exposed cysteine residues C469 and C473 (Fig. 4a,b). While we were able to readily generate strain His_8_-*PFKELCH13^C469S^* (Fig. S3b), three SLI attempts to generate His_8_-*PFKELCH13^C473S^* were unsuccessful (two of which were again performed in parallel to successful SLI experiments). Taking into account that only a single surface-exposed oxygen atom is replaced in His_8_-*PFKELCH13^C473S^*, we speculate that the thiol group of residue C473 might be essential. In contrast, serine mutants His_8_-*PFKELCH13^C469S^*, His_8_-*PFKELCH13^C532S^*, and His_8_-*PFKELCH13^C580S^* showed no growth defect (Fig. 4d).

In summary, strain NF54K13^C580Y^ has a normal susceptibility to common redox agents. Not the loss of the thiol group of C580 but the specific replacement by tyrosine increases the artemisinin susceptibility of *P. falciparum*. However, a Keap1-like redox function of PfKelch13 cannot be excluded at the current stage because of three unsuccessful attempts to replace the thiol group of surface-exposed residue C473.

### Characterization of recombinant PfKelch13 and antibody generation

To the best of our knowledge, no protocol for the production of recombinant PfKelch13 has been published yet despite the deposition of PDB entries 4ZGC and 4YY8 by Jiang *et al.* and the Structural Genomics Consortium (SGC) (DOI: 10.2210/pdb4zgc/pdb). To study the effects of common mutations as well as the protein properties of PfKelch13 in more detail, we produced recombinant full-length PfKelch13 as well as truncated PfKelch13^337–726^ that comprises the BTB domain and the Kelch β-propeller domain (Fig. 5a). Recombinant full-length PfKelch or PfKelch13^337-726^ produced in *Escherichia coli* was insoluble despite numerous parameter variations (including alternative strains, temperatures, IPTG- or autoinduction, N-terminal His_6_-, GST- or MBP-tags or the systematic replacement of cysteine residues), indicating intrinsic protein folding problems most likely because of the β-propeller domain (Table S1). We also tested the production of His_8_-tagged PfKelch13 and PfKelch13^337-726^ in *Pichia pastoris* GS115 and the *Leishmania tarentolae*-based LEXSY system but did not obtain soluble recombinant protein form these eukaryotes. Nevertheless, we established a solubilization and purification protocol for recombinant MAH_6_VGT-tagged PfKelch13^336-726^ from inclusion bodies in *E. coli* to generate and affinity-purify an antibody against PfKelch13 (Fig. S4a-c). Western blot analyses using this antibody revealed a specific detection of endogenous PfKelch13 at ∼85 kDa in *P. falciparum* lysates. Furthermore, we coupled the antibody to CNBr-activated sepharose and pulled-down denatured PfKelch13 and His_8_-PfKelch13^wt^ from parasite lysates of strains 3D7 and His_8_-*PFKELCH13^wt^*, respectively (Fig. S4d). The eluate containing His_8_-PfKelch13^wt^ was probed with an antibody against the His-tag (Fig. S4d) and was also analyzed by mass spectrometry (as outlined below), confirming the specifity of the antibody against PfKelch13 and the successful pull-down. Since disulfide bonds are maintained under denaturing conditions, the established protocol can be applied in future studies to identify potential redox interaction partners using wild-type PfKelch13 and cysteine mutants.

**Figure 5.**
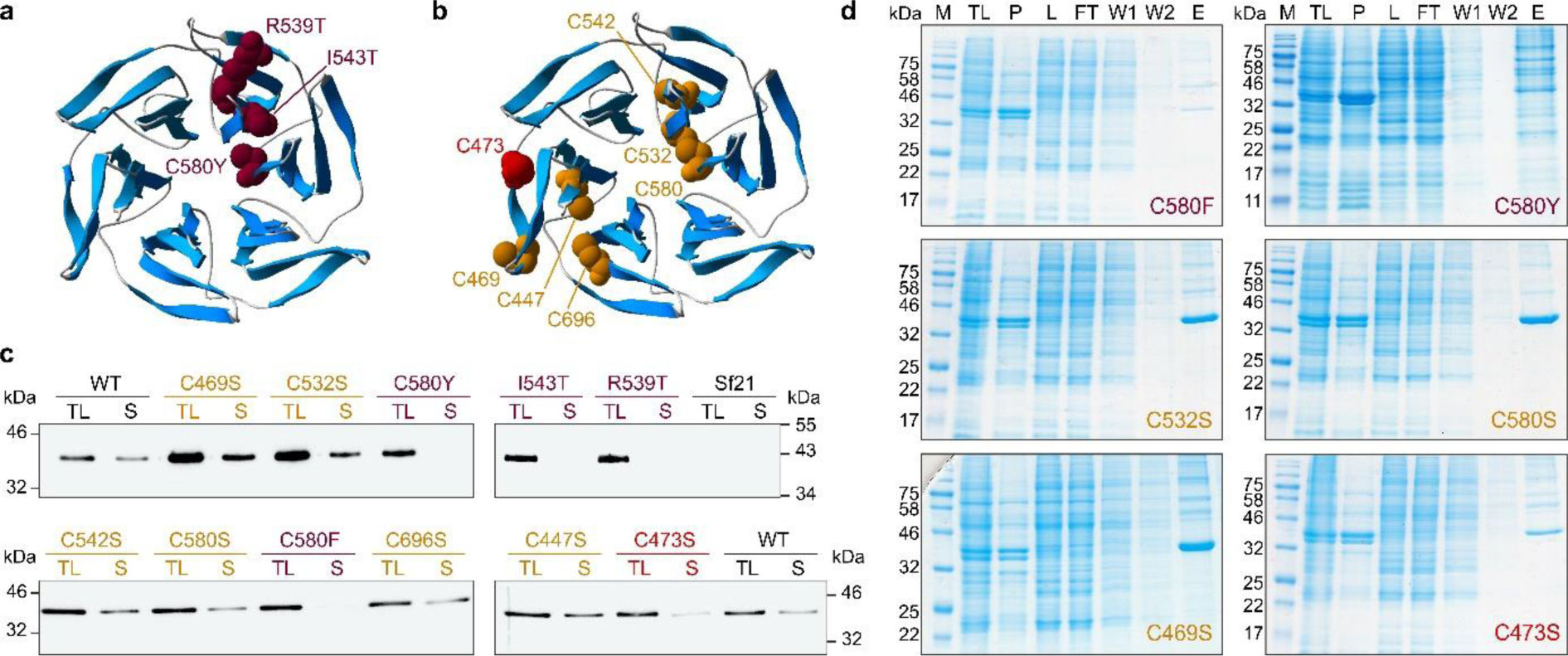
Solubility of recombinant PfKelch13^337-726^ mutants. (**a**) Overview of three residues in the β-propeller of PfKelch13 (PDB entry 4YY8) that are frequently mutated in *P. falciparum* field isolates and that are linked to a decreased artemisinin susceptibility. (b) Overview of the seven cysteine residues in the β-propeller of PfKelch13 including disulfide-bridged C532 and C580 (PDB entry 4YY8). (**c**) Solubility screen of wild-type (WT) and mutant MH_8_SR-tagged PfKelch13^337-726^ in Sf21 cells. Cells were lysed by three freeze-thaw cycles and equal volumes of the total lysate (TL) and the soluble fraction (S) were separated by SDS PAGE and analyzed by western blotting against the His-tag. Lysates from SF21 cells without PfKelch13^337-726^ served as a negative control. **d**) Representative purifications of mutant PfKelch13^337-726^ from panel c. Cells were lysed by three freeze-thaw cycles. The total lysate (TL) was centrifuged, yielding insoluble pellet (P) and supernatant (L), which was loaded on a Ni-NTA agarose column. The flow-through (FT) was discarded, the resin was washed (W) and the bound protein eluted (E) with imidazole.

Next, we produced and purified recombinant MH_8_SR-tagged PfKelch13^337-726^ from *Spodoptera frugiperda* Sf21 cells (Fig. S5a). The yield was around 6 mg of pure soluble protein per liter of culture. SDS-PAGE analysis showed an apparent molecular mass of 43 kDa in accordance with the calculated molecular mass of 45.8 kDa. Mass spectrometry with a sequence coverage of 74% confirmed that the purified protein was the expected product and that the C-terminus was intact (Fig. S5b). Circular dichroism (CD) spectroscopy revealed a high percentage of regular secondary structure elements, with α-helices and β-sheets amounting to about 24% each (Fig. S5c). These values deviate from the values of 10% α-helix and 34% β-sheet derived from the crystal structure of the protein (PDB 4YY8), which might suggest a higher percentage of α-helices in solution as compared with the crystal. Interestingly, CD spectroscopy of the structurally similar protein Keap1 also suggested a flexible structure^24^ and a higher α-helix content in solution than in the crystal structure^34^. We also probed the thermostability of the folded state of the protein by monitoring its secondary structure composition as a function of temperature with the aid of CD spectroscopy (Fig. S5d). Irrespective of the wavelength, a sharp transition was observed within a narrow temperature interval around 55°C. Above this temperature, the CD signal decreased in magnitude and virtually disappeared even at those wavelengths where an unfolded polypeptide chain would manifest in a pronouncedly negative ellipticity (e.g., 192 nm and 198 nm). This finding is in agreement with the observed protein precipitation at temperatures above 50°C. Correct protein folding was corroborated independently by measuring the accessibility of cysteine residues. In accordance with the predicted accessibility of two out of seven cysteine residues (Fig. 4a), only two (1.8 ± 0.2) cysteine residues were found to be accessible to Ellman’s reagent under native conditions, whereas all seven (6.8 ± 0.4) cysteine residues became accessible under denaturing conditions (Fig. S5d). Gel filtration chromatography revealed the presence of multiple oligomeric states (Fig. S5e). We detected either monomeric recombinant PfKelch13^337-726^ in 50 mM sodium phosphate buffer without salt or the interconversion of monomeric, dimeric, and tetrameric protein in the same buffer containg 300 mM NaCl, suggesting a dynamic equilibrium between these three species.

In summary, we generated a specific antibody and established a denaturing pull-down assay for PfKelch13, developed a protocol for the purification of soluble recombinant PfKelch13^337-726^ from Sf21 cells, confirmed that two of the seven cysteine residues are surface exposed, and detected an unexpectedly flexible quaternary structure.

### Potential post-translational modification of PfKelch13

Even though recombinant His-tagged full-length PfKelch13 produced in *E. coli* and Sf21 cells was insoluble, lysates allowed us to compare the electrophoretic mobility with PfKelch13 from *P. falciparum* by SDS-PAGE and western blot analysis. While unmodified PfKelch13 from strain 3D7 was detected at 95 kDa, His-tagged full-length PfKelch13 from strain His_8_-*PFKELCH13^wt^* had an apparent molecular mass of approximately 97 kDa (Fig. S6a). The mass difference between both proteins was in accordance with the modification of the N-terminus of His_8_-PfKelch13^wt^ due to the ribosomal skipping and His-tag, however, both proteins ran about 10 to 12 kDa higher than expected. The mass shift was detected in blots that were probed with our anti-PfKelch13 antibody and in blots that were probed with a commercial antibody against the His-tag. Recombinant His-tagged PfKelch13 from *E. coli* or Sf21 cells was detected around 85 kDa in accordance with the calculated molecular mass (Fig. S6a). These controls disproved an inherent abnormal mobility of PfKelch13 and confirmed the mass shift of PfKelch13 in *P. falciparum*. A second band with variable intensity around the expected molecular mass of 85 kDa was also detected in *P. falciparum* lysates that were probed with the anti-His antibody (Fig. 2b) or the anti PfKelch13 antibody (Fig. S6a). Although both PfKelch13 species were regulated in the knockdown experiments (Fig. 2b), the intensity of the signal at 85 kDa and the ratio between the upper and lower band differed among experiments, suggesting a dynamic interconversion of the protein species during blood-stage development.

The modified gene architectures of His_8_-*PFKELCH13-glmS* (Fig. 1a) and His_8_-*PFKELCH13^wt^* (Fig. S3a) refuted an alternative translation initiation site that is encoded at the 5’ end of *PFKELCH13*. A putative translational read-through until the next stop codon would have increased the mass by just 1.8 kDa. Thus, a post-translational modification is the most plausible explanation for the mobility shift of PfKelch13 in *P. falciparum*. Since pull-down experiments worked well with our anti-PfKelch13 antibody (Fig. S4d), we excised the band corresponding to PfKelch13 and analyzed it by mass spectrometry (Fig. S6b). Although highest signal intensities corresponded to peptides of PfKelch13, we neither detected proteins nor post-translational modifications that could explain the altered electrophoretic mobility. In summary, we identified two different forms of PfKelch13 with distinct electrophoretic mobilities around 85 and 95 kDa in *P. falciparum*, suggesting an unidentified post-translational modification.

### Protein folding rather than redox properties correlates with artemisinin susceptibility

Next, we produced single point mutants of MH_8_SR-tagged PfKelch13^337-726^ in Sf21 cells to analyze the altered protein properties. Mutations R539T, I543T and C580Y cause a decreased susceptilibity against artemisinin in the field and in cell culture^7,9,15,31^ and are all within the propeller domain (Fig. 5a). Furthermore, we systematically replaced the seven cysteine residues (Fig. 5b). Western blot analysis of the Sf21 lysates with the according recombinant proteins revealed that each of the field mutations had a strong destabilizing effect, resulting in completely insoluble recombinant PfKelch13^337-726^ (Fig. 5c). This was also observed for the replacement C580F, whereas single replacements C447S, C469S, C532S, C542S, C580S, and C696S had no or only moderate effects on the protein solubility. An anomaly with intermediate solubility was observed for the replacement C473S. Protein purifications confirmed the different solubilities (Fig. 5d). In summary, not the loss of the disulfide bond between residues C532 and C580 but the C580Y or C580F replacement led to insoluble PfKelch13^337-726^. Impaired protein folding of recombinant PfKelch13^337-726^ correlates with the observed decreased artemisinin susceptibility of field mutants, suggesting that protein folding is the central common parameter for mutant selection.

## Discussion

The relevance of PfKelch13 as an artemisinin susceptibility factor can be viewed from two different perspectives, one with a focus on the chemical properties and mode of action of artemisinin and one with a focus on the properties and structure-function relationships of PfKelch13. The endoperoxide group of artemisinin is a prerequisite for its antimalarial activity^35^. While artemisinin itself seems to be not a peroxidase substrate^33^, heme-dependent one electron reduction of the endoperoxide bond results in a reactive radical that was shown to alkylate heme in cell culture and mice^36, 37^ as well as proteins *in vitro* or in cell culture^36, 38, 39^. Furthermore, artemisinin-dependent inhibition of hemozoin formation was suggested to cause oxidative stress^40^. Thus, artemisinin might either directly or indirectly modify C580 or another cysteine residue in PfKelch13. A direct alkylation of PfKelch13 by artemisinin seems less likely considering the absent co-localization of PfKelch13, heme, and highly reactive artemisinin radicals. Modifications of specific cysteine residues of the vertebrate homologue Keap1 by hydrogen peroxide or alkylating agents is a key event in signal transduction that leads to the transcription of numerous genes and an increased biosynthesis of glutathione^21^. Rodent and human Keap1 have 25 and 27 cysteine residues, respectively. Two of these residues in the intervening region were shown to act as a sensor and to react with electrophiles^22, 41–43^. Another regulatory cysteine that is activated by a different set of electrophiles was found in the BTB domain, suggesting a highly complex integration of diverse signals^22, 24, 42, 43^. The homology between Keap1 and PfKelch13 leads to the hypothesis that PfKelch13 also acts as a redox sensor. Mutations in *PFKELCH13* were indeed shown to correlate with an increased abundance of γ-glutamylcysteine and glutathione^44^. However, PfKelch13 lacks the intervening region of Keap1, all its seven cysteine residues are located in the β-propeller domain, and serine replacements for C469, C532 or C580 of PfKelch13 had no effect on the artemisinin susceptibility in cell culture. Thus, a potential redox regulation of these residues is irrelevant for the altered artemisinin susceptibility. If PfKelch13 undergoes an essential redox regulation and/or an artemisinin-dependent redox regulation, it has to depend on the thiol group of residue C473, which was refractory to replacement in *P. falciparum* blood stages and which also affected the folding properties of recombinant PfKelch13.

We provide evidence that PfKelch13 variants in the field have an altered protein stability (Fig. 6). We established a protocol for the production of recombinant PfKelch13 and showed that each of the common mutations R539T, I543T or C580Y destabilizes the protein and decreases its solubility in accordance with molecular dynamics simulations^10^. While mutations C580S or C532S had no effect on protein folding, mutations R539T, I543T, C580Y or C580F all resulted in misfolded insoluble recombinant proteins. Thus, protein stability and abundance link the previously detected decreased or normal artemisinin susceptibility following protein mislocalization of wild-type PfKelch13 or overexpression of *PFKELCH13^C580Y^*, respectively^15–17^. A correlation between protein abundance and artemisinin susceptibility was previously shown by quantitative mass spectrometry^17, 44^. We now confirmed the link between protein abundance and artemisinin susceptibility using a titratable knockdown system (Fig. 6). The knockdown resulted in arrested ring-stage parasites during the second intraerythrocytic cycle. This result is in accordance with previous mislocalization studies showing that PfKelch13 is not essential for schizont development^16^. The developmental arrest was shown to correlate with a decreased PfKelch13-dependent endocytotic uptake and proteolytic degradation of hemoglobin in early ring-stage parasites^16, 17, 44^, which is thought to prevent the heme-dependent activation of artemisinin and to give the parasite time to survive the peak concentration of the rather labile drug^6,40, 45^. Our knockdown parasites reached ring-stage survival percentages slightly above 40%. This value might indicate a physiological threshold because knockdown parasites died at higher GlcN concentrations and similar maximum survival percentages were reported for RSAs with dihydroartemisinin and mutants R539T and I543T in other genetic backgrounds^9^. The degree of misfolding of common β-propeller domain mutants in *P. falciparum* remains to be analyzed. However, misfolding of the β-propeller domain or decreasing the amount of PfKelch to less than 10% was still compatible with the essential function of PfKelch13, which was shown to require the N-terminal apicomplexan-specific region^16^. Lower PfKelch13 abundances presumably limit the endocytotic uptake of nutrients to such a degree that the parasite cannot develop any further.

**Figure 6.**
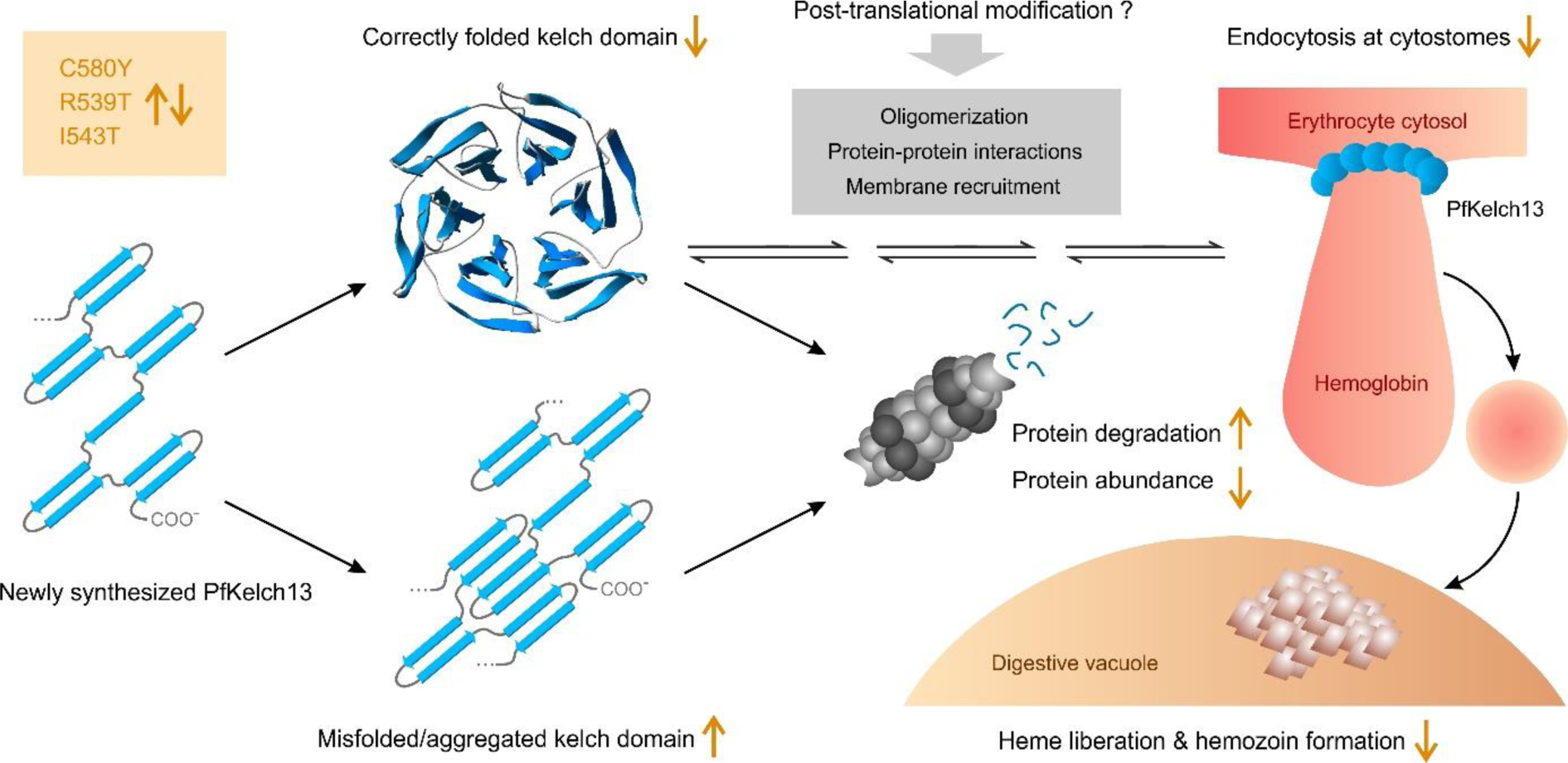
Updated model for the decreased artemisinin susceptibility of strains with mutant or down-regulated PfKelch13. Common field mutations destabilize the kelch domain, resulting in a decreased protein abundance, endocytosis, and liberation of heme, which activates artemisinin. This phenotype can be mimicked in parasites with down-regulated PfKelch13.

Our comparative western blot analyses with recombinant and endogenously tagged as well as untagged PfKelch13 revealed two different forms of PfKelch13 with distinct electrophoretic mobilities around 85 and 95 kDa. It is unlikely that the double band and mass shift were caused by limited proteolysis as the C-terminus of PfKelch13 is buried according to PDB entry 4YY8 and the N-terminal His-tag was detected for both recombinant and endogenously tagged PfKelch13. Since we can also exclude an alternative translation initiation site or a translational read-through as potential causes for the mass shift, we suggest that a major fraction of PfKelch13 undergoes a post-translational modification. The potential modification remains to be identified. Attachment of a protein, such as SUMO or ubiquitin, appears to be the most likely modification, not only because of the mass shift but also because PfKelch13 lacks common peptide motifs for lipid modifications (and multiple phosphorylations or acetylations would rather increase the electrophoretic mobility due to additional negative charges or the loss of positive charges, respectively). It also remains to be determined whether a post-translational modification regulates (i) the alternative oligomeric states that were observed for recombinant PfKelch13 *in vitro*, (ii) protein-protein interactions, and/or (iii) the membrane recruitment of PfKelch13 to different subcellular compartments, including the doughnut-shaped structures at cytostomes^16–18^ (Fig. 6).

In conclusion, we showed that down-regulation of PfKelch13 results in ring-stage survival rates of up to 40% and that common field mutations have a destabilizing effect on the folding properties of PfKelch13. We also showed that PfKelch13 exists in at least two different forms and established a protocol for the production of recombinant PfKelch13^337–726^. The recombinant protein adopts alternative oligomeric states and two of its seven cysteine residues react with DTNB. While the thiol group of C473 of PfKelch13 appears to be essential for blood-stage development, the redox state of C580 and C532 is irrelevant for the PfKelch13-dependent artemisinin susceptibility.

## Online methods

### Cloning of constructs

Primers were purchased from Metabion and are listed in Supplementary Tables S2 – S4. SLI constructs were generated using vector pSLI-N-GFP-2xFKBP-loxP(K13)^15^ (addgene plasmid #85792). Insert GFP-2xFKBP-K13 was excised with *Nhe*I and *Xho*I. Recodonized *PFKELCH13* was amplified by PCR using primers P65 and P69 and vector pSLI-N-GFP-2xFKBP-loxP(K13) as a template. The PCR product was digested with *Sal*I and *Xho*I. Primers P60 and P61 that encode the His_8_-tag were annealed, yielding cohesive ends for *Nhe*I and *Sal*I. The annealed primers were ligated with the digested vector and PCR product, yielding pSLI-His_8_-*PFKELCH13*. To introduce mutations in recodonized *PFKELCH13*, the gene was first subcloned into plasmid pET45b using restriction sites *Avr*II and *Xho*I, followed by site-directed mutagenesis with primers P71 – P76 or P84 – P87, and recloning into pSLI-N-GFP-2xFKBP-loxP(K13) using primers P65 and P69. Primers P88 and P89 were used to amplify the *glmS* and *M9* sequences from plasmids pCR 2.1 TOPO-*mCherry*-*glmS* and pCR 2.1 TOPO-*mCherry*-*M9*, respectively^46^. The loxP site from pSLI-His_8_-*PFKELCH13* was subsequently excised and replaced with the *glmS* or *M9* sequence using *Stu*I and *Xho*I, yielding pSLI-His_8_-*PFKELCH13*-*glmS* and pSLI-His_8_-*PFKELCH13*-*M9*.

Plasmid pET45b-*PFKELCH13* with synthetic *PFKELCH13* that was codon-optimized for the expression in *E. coli* (Fig. S7) was purchased from GenScript and was used as a template to PCR-amplify truncated *PFKELCH13^337-726^* with primers P1 and P2. The PCR product was cloned into the *Kpn*I and *Avr*II restriction sites of pET45b, resulting in construct pET45b-*PFKELCH13^337-726^*. Plasmids pET45b-*PFKELCH13* and pET45b-*PFKELCH13^337-726^* served as templates for site-directed mutagenesis with primers P6 – P23 to generate serine mutants for each of the seven cysteine residues as well as mutants C580Y and C580F. Synthetic *PFKELCH13* and *PFKELCH13^337-726^* from plasmid pET45b were also PCR-amplified and subcloned into the *Bam*HI and *Hin*dIII restriction sites of pQE30 (using primers P3 – P5), the *Bam*HI and *Not*I restriction sites of pGEX-4T-1 (using primers P3, P4 and P30), the *Nde*I and *Not*I restriction sites of pMAL-c5X (using primers P30, P32 and P33), the *Eco*RI and *Not*I restriction sites of pBLHIS-SX (using primers P30, P37 and P39), and the *Xba*I and *Not*I restriction sites of pLEXSY-IE-blecherry4 (using primers P30, P36 and P38).

Constructs for expression in Sf21 cells were based on plasmid pCoofy41 (addgene plasmid #55184)^47^, which was digested with *Bam*HI and *Hin*dIII. *PFKELCH13^337-726^* was amplified by PCR from wild-type and mutant pET45b-*PFKELCH13* using primers P26 and P4. The PCR products were digested with *Xba*I and *Hin*dIII. Primers P50 and P51 that encode the His_8_-tag were annealed, yielding cohesive ends for *Bam*HI and *Xba*I. The annealed primers were ligated with the digested vector and PCR product, yielding wild-type or cysteine mutant-encoding pCoofy-His_8_-*PFKELCH13^337-726^*. Two additional mutant versions (R539T and I543T) were generated by site-directed mutagenesis with primers P95 – P98 using pCoofy-His_8_-*PFKELCH13^337-726^* as a template. Construct pCoofy-His_8_-*PFKELCH13* encoding full-length PfKelch13 was generated analogously with primers P52 and P4. All constructs were confirmed by Sanger sequencing (SEQ-IT GmbH & Co. KG).

1. *P. falciparum* culture, transfection and selection.
2. *P. falciparum* strains were cultured according to Trager and Jensen^48^ with slight modifications^49^ at 37°C, 5% O_2_, 5% CO_2_, and 90% humidity in fresh human A^+^ erythrocytes at a hematocrit of 3.6% in complete RPMI-1640 medium containing 0.45% AlbuMAX II, 0.2 mM hypoxanthine, and 5 µg/mL gentamicin. Standard 14 mL cultures were maintained in vented petri dishes.

Transfections were performed with pre-loaded uninfected erythrocytes that were subsequently infected with *P. falciparum*^50, 51^. Parasites were synchronized with 5% sorbitol^52^ one day before transfection. To sterilize the plasmid DNA, 100 µg of a DNA from a midi-preparation was acidified with 0.1 volumes of buffer containing 3 M sodium acetate/acetic acid, pH 4.7 and precipitated with 2.5 volumes of absolute ethanol at -20°C over night. After centrifugation at 20000 × *g* for 30 min at 4°C, the DNA was washed with 500 µL of ice-cold 70% ethanol. The supernatant was discarded under sterile conditions and the DNA was dried for 10 min before the transfection. Fresh erythrocytes were washed twice with 3.5 volumes of cytomix containing 120 mM KCl, 5 mM MgCl_2_, 0.15 mM CaCl_2_, 2 mM EGTA, 10 mM K_2_HPO_4_, 10 mM KH_2_PO_4_, 25 mM HEPES/KOH, pH 7.6 at room temperature. An aliquot of 400 µL washed erythrocytes was mixed with 100 µg of sterile DNA in 400 µL cytomix, transferred to two electroporation cuvettes, incubated on ice for 5 min, and electroporated using program U-033 of the Nucleofector 2b (Lonza). Following electroporation, both cuvettes were immediately incubated for 5 min on ice. Pre-loaded erythrocytes were combined after rinsing each cuvette with 4 mL complete RPMI-1640 medium, centrifuged at 300 × *g* for 5 min, and resuspended together with 100 µL infected erythrocytes with 1% parasitemia in complete RPMI-1640 medium. Cultures were washed the next day and supplemented with 5 nM of the antifolate WR99210 in a new petri dish. Drug and medium were changed daily during the first week. Afterwards, WR99210 and medium were renewed every other day, and 50 µL of fresh erythrocytes were added once a week until parasites were detected by light microscopy in Giemsa-stained blood smears 3–5 weeks post-transfection.

SLI was performed with successfully transfected and selected parasites containing pSLI plasmids^15^. Cultures were grown to a parasitemia of 5% and supplemented with 0.9 µM DSM1. Drug and medium were renewed daily. Usually, parasites disappeared after 4 days and reemerged after around 2 weeks. Successful integration was confirmed by genotyping PCRs with extracted genomic DNA^53^ and western blot analysis against the His_8_-tag following parasite isolation by saponin treatment^54^.

### Growth curve analyses, IC_50_ measurements and RSAs

A 1.16 mL stock solution of 400 mM GlCN was prepared freshly by neutralizing 100 mg GlcN*HCl (Sigma) with 232 µL 2 M NaOH and adding 928 µL RPMI-1640 medium. Asynchronous cultures of strains His_8_-*PFKELCH-glmS* and His_8_-*PFKELCH-M9* were set to a parasitemia of 0.1%, split into four and treated with either 0, 0.3, 1.0, 2.0, 5.0 or 10 mM GlcN for five days. The medium with the corresponding GlcN concentration was replaced once a day. Giemsa-stained blood smears were analyzed by light microscopy. At least 2000 erythrocytes were counted to determine the parasitemia for each day. The data from at least three independent growth experiments was analyzed in SigmaPlot13.

IC_50_ measurements were performed in a SYBR green I plate-reader assay as described previously^32, 55^ with slight modifications. Parasite cultures were synchronized twice with sorbitol. Stock solutions of 0.49 µM artesunate, 39 mM tBuOOH, 49 mM diamide, 122 mM DTT, and 97 µM DSM1 were freshly prepared in albumax-free RPMI-1640 medium containing 0.2 mM hypoxanthine and 5 µg/mL gentamicin, and were subsequently filter-sterilized. Early ring-stage parasites in 100 µL complete medium at a parasitemia of 0.3% and a hematocrit of 1.5% were treated in 96-well plates with 0.01 – 81 nM artesunate, 1.0 µM – 6.5 mM tBuOOH, 1.2 µM – 8.1 mM diamide, 3.0 µM – 20 mM DTT, or 2.5 nM – 16 µM DSM1, incubated for 140 h, and then stored at -80°C. Uninfected erythrocytes as well as untreated parasites served as controls. Cells were thawed and lysed in 100 µL lysis buffer (20 mM Tris-HCl, pH 7.5, 5 mM EDTA, 0.08% (v/v) Triton X-100, 0.008% (w/v) saponin) that was supplemented with 0.12 µL/mL 10000 × SYBR green I solution, and the fluorescence was analyzed using a ClarioStar plate reader (BMG Labtech) with an excitation at 485 nm and an emission at 535 nm. The gain and optimal measuring height were internally adjusted to the highest expected fluorescence intensities for the parasite cultures without drug. The fluorescence intensities of the uninfected erythrocytes with the corresponding drug concentrations were substracted from the intensities of the parasite cultures. The resulting fluorescence intensities were normalized with regard to the maximum intensities from untreated controls, plotted against the corresponding drug concentrations, and fitted to a sigmoidal dose-response curve using a four-parameter Hill function in SigmaPlot13.

RSAs were performed as described previously^4,33^ with slight modifications. Early ring-stage cultures were highly synchronized by three consecutive sorbitol treatments with a time difference of 45 h. The synchronized culture was split in two and set to a parasitemia of 1% and a hematocrit of 2%. One 14 mL culture was treated with 700 nM artesunate. The other culture was treated with the same DMSO concentration and served as a control to calculate the survival percentage. After 6 h, the cultures were washed four times with 5 mL complete RPMI-1640 medium, resuspended in 14 mL complete RPMI-1640 medium, and cultured for another 66 h. The parasitemia was determined from Giemsa-stained blood smears.

The formation of hemozoin was analyzed for highly synchronized strains His_8_-*PFKELCH-glmS* and His_8_-*PFKELCH-M9*. One cycle before the assay, parasites were pre-synchronized with sorbitol and incubated with 50 units of heparin to inhibit invasion. When the majority of parasites presented as segmenters, heparin was washed off and invasion was allowed for 2 h under constant agitation at 80 rpm. All remaining late-stage parasites were removed by sorbitol treatment and cultures were treated with 1 mM GlcN. Medium with GlcN was replaced once per day and thin blood smears were taken at 18–28 h hpi every 2 hours for the first cycle. After the 28 hpi timepoint of the first cycle, cultures were again treated with 50 units of heparin. When the majority of parasites presented as segmenters, invasion was allowed for 1 h at 80 rpm. All remaining late-stage parasites were removed by sorbitol treatment. For the second cycle, thin blood smears were taken at 16–32 hpi every 2 hours. Parasites were examined for hemozoin by light microscopy of thin blood smears that were stained with Hemacolor rapid staining of blood smear solution (Sigma) using a BZ-X800 microscope (Keyence).

### Recombinant PfKelch13 and antibody generation

To produce His_8_-tagged PfKelch13 and PfKelch13^337-726^ in *E. coli*, the full-length and truncated constructs in pET45b, pQE30, pGEX-4T-1 and pMAL-c5X were tested using the strains and conditions listed in supplementary Table 1. Recombinant PfKelch13^337-726^ for the generation of antibodies was produced in *E. coli* SHuffle T7 Express cells (NEB) containing pET45b-*PFKELCH13^337-726^*. Cells were grown in an Innova 44R shaker at 170 rpm at 30°C and induced with 0.5 mM isopropyl β-D-1-thiogalactopyranoside (IPTG) at an opitical density (OD) of 0.5. After 4 h, cultures were incubated for 10 min in an ice-water bath and centrifuged in a Beckman JS-4.2 rotor at 4000 × *g* for 15 min at 4°C. The cell pellet from 1 L culture was resuspended in 10 mL ice-cold purification buffer (300 mM NaCl, 50 mM Na_x_H_y_PO_4_, pH 8.0) and stored at -20°C. The thawed suspension was supplemented with 10 mg lysozyme and a spatula tip of DNase, stirred on ice for 1 h, sonicated on ice and centrifuged at 10000 × *g* for 30 min at 4°C. The pellet containing the inclusion bodies was washed twice at room temperature with 10 mL washing buffer (300 mM NaCl, 2% (w/v) Triton X-100, 2 M urea, 50 mM Tris-HCl, pH 8.0) followed by sonication and solubilization in 10 mL solubilization buffer (300 mM NaCl, 2% (w/v) SDS, 8 M urea, 50 mM Tris-HCl, pH 8.0). Excess SDS was precipitated at 4°C overnight followed by centrifugation at 10000 × *g* for 30 min at 4°C. The supernatant was loaded on a 500 µL Ni-NTA agarose column (Qiagen), washed with 10 mL solubilization buffer without SDS and eluted with 1.5 mL of the same buffer containing 200 mM imidazole, pH 8.0. Eluted PfKelch13^337-726^ was washed and concentrated to 60 µM in solubilization buffer without SDS using a Amicon Ultra-15, PLGC Ultracel-PL Membran, 10 kDa unit (Merck) and heated at 95°C for 10 min in the presence of 40 mM DTT to reduce all cysteines. The solubilized denatured protein from *E. coli* was used for commercial immunization of rabbits (Pineda Antikörper Service, Berlin) and was also coupled to CNBr-activated sepharose^56^ for the purification of PfKelch13 antibodies from rabbit serum by affinity-chromatography.

Recombinant His_8_-tagged PfKelch13 and PfKelch13^337-726^ were produced in Sf21 cells that were routinely cultured in Sf900^TM^ III SFM medium (Thermo) in an Innova 44R shaker at 120 rpm at 27°C. The *PFKELCH13*-encoding pCoofy constructs were used for transposition into *E. coli* DH10EMBacY cells (Geneva Biotech) and bacmid DNA was isolated from 3 mL overnight cultures. For the transfection, a solution containing 10 µg of bacmid DNA and 50 µL of medium were mixed with a second solution comprising 5 µL of X-tremeGENE HP DNA transfection reagent (Merck) and 50 µL medium. The transfection mixture was incubated for 10 min at room temperature and then added dropwise to 0.6 × 10^6^ adherent Sf21 cells in 2 mL medium in a 6-well Corning plate. After 5 h of incubation at 27°C, 1 mL of medium was added to each well and the plate was incubated for 3 days at 27°C. The cell culture medium containing the V_0_ baculovirus stock was harvested and 2 mL of V_0_ was used to infect 25 mL Sf21 cells at a density of 0.6 × 10^6^ cells/mL in a 125 mL Erlenmeyer flask to generate the V_1_ baculovirus stock. Once the cells stopped proliferating, they were kept in culture for an additional 24 h and then the baculovirus (V_1_ stock) was harvested by centrifugation (10 min, 550 × *g*). For recombinant protein production, 5 mL of V_1_ was added to 1 L of Sf21 culture at a density of 10^6^ cells/mL. After 72 h, cells were harvested (10 min, 650 × *g*) and stored at -20°C. The cell pellet was resuspended in 30 mL ice-cold lysis buffer (20 mM imidazole, 300 mM NaCl, 10 mM MgCl_2_, 1 spatula tip DNase, 1 tablet cOmplete EDTA-free (Roche), 50 mM Na_x_H_y_PO_4_, pH 8.0) and divided into four samples. Cells were disrupted by three freeze-thaw cycles using liquid nitrogen and centrifuged at 10000 × *g* for 30 min at 4°C. The combined supernatants were loaded on a 0.8 mL Ni-NTA agarose column that was equilibrated with 10 mL buffer containing 20 mM imidazole, 300 mM NaCl, 50 mM Na_x_H_y_PO_4_, pH 8.0. The column was washed with 10 mL ice-cold washing buffer (50 mM imidazole, 300 mM NaCl, 50 mM Na_x_H_y_PO_4_, pH 8.0) and the protein was eluted with buffer containing 250 mM imidazole, 300 mM NaCl, 50 mM Na_x_H_y_PO_4_, pH 8.0.

### Gel filtration chromatography, Ellman’s assay and CD spectroscopy

The oligomerization state of recombinant PfKelch13^337-726^ freshly purified from Sf21 cells was analyzed at 10°C on a Superdex 200 Increase 10/300 GL column connected to an Äkta explorer system using 50 mM Na_x_H_y_PO_4_, pH 8.0 with or without 300 mM NaCl as a running buffer.

The number of accessible cysteines in PfKelch13^337-726^ was determined with Ellman’s reagent 5,5’-dithiobis-(2-nitrobenzoic acid) (DTNB)^57^. Freshly purified recombinant PfKelch13^337-726^ was reduced with 5 mM DTT for 30 min on ice. Imidazole and excess DTT were removed using a PD-10 desalting column and phosphate buffer (300 mM NaCl, 50 mM Na_x_H_y_PO_4_, pH 8.0). The protein was diluted with phosphate buffer to a concentration of 10 µM and 30 µM using the calculated molar extinction coefficient of ε_280nm_ = 59.82 mM^-1^cm^-1^. Samples were mixed with equal volumes of 1.2 mM DTNB in either phosphate buffer or denaturing buffer (2% (w/v) SDS, 300 mM NaCl, 50 mM Na_x_H_y_PO_4_, pH 8.0). After 5 min incubation at room temperature, the absorbance was measured at 412 nm using a JASCO V-650 UV/Vis spectrophotometer. Calibration curves with 0 – 40 µM DTT were generated in parallel for both buffers and were used to calculate the concentration of accessible thiols under native and denaturing conditions from three independent protein purification experiments.

For CD spectroscopy, freshly purified recombinant PfKelch13^337-726^ was concentrated and the buffer exchanged with CD buffer (100 mM NaF, 50 mM Na_x_H_y_PO_4_, pH 7.4) in an Amicon Ultra-15, PLGC Ultracel-PL Membran, 10 kDa unit. CD spectra were recorded at 20°C using a thermostatted Chirascan CD spectrometer (Applied Photophysics). The proportions of secondary structure elements were estimated from the spectra using the K2D2 software^58^. The contents of secondary structure elements in the crystal structure of PfKelch13^337-726^ (PDB entry 4YY8) were calculated using the webserver 2StrucCompare.

### Immunoprecipitation of PfKelch13

For immunoprecipitation of PfKelch13 from *P. falciparum*, four 14 ml blood-stage cultures with a parasitemia of ∼10% trophozoites were combined and harvested by centrifugation at 300 × *g* for 5 min. Following saponin treatment^54^, parasites were resuspended in 250 µL lysis buffer (150 mM NaCl, 2% (w/v) SDS, 1% (v/v) Triton X-100, 10 mM Tris-HCl, pH 7.5) containing cOmplete EDTA-free protease inhibitor (Roche) and frozen in liquid nitrogen. After thawing, the lysate was cleared by centrifugation at 16000 × *g* for 10 min at room temperature. The supernatant was diluted 1:20 with buffer containing 150 mM NaCl, 10 mM Tris-HCl, pH 7.5 and incubated at 4°C overnight rotating end over end with PfKelch13 antibody that was coupled to 100 µL CNBr-activated sepharose. The sepharose was centrifuged at 500 × *g* for 5 min at 4°C and washed three times with 1 mL 150 mM NaCl, 10 mM Tris-HCl, pH 7.5. Bound proteins were eluted by adding 100 µL of 2x Laemmli buffer and heating at 95°C for 5 min. Samples were subjected to SDS-PAGE and western blot analysis.

### Data availability

All relevant data are included in the manuscript or its supplementary information and are available from the authors upon request.

## Acknowledgments

This work was funded by the Deutsche Forschungsgemeinschaft (DFG, German Research Foundation) through the priority program SPP 1710 (grant DE 1431/8-2 to M.D.) and the SFB 1129 (project number 25240245660 to M.G.) as well as the Baden-Württemberg Foundation (ref: 1.16101.17 to M.G.). We thank Frederik Sommer and Michael Schroda for the mass spectrometry analysis of recombinant PfKelch13^337-726^ from Sf21 cells, Markus Räschle for the mass spectrometry analysis of purified His_8_-Kelch13 from *P. falciparum*, Simone Eggert and Stefan Kins for their support with PfKelch13 expression trials in *P. pastoris*, and Tobias Spielmann for SLI plasmids and helpful discussions.

## Author Contributions

M.D. and R.S. conceived the study design. R.S. conducted and M.D. supervised all experiments unless otherwise indicated. The hemozoin analysis was conceived and supervised by M.G. and performed by S.Kl., S.Ke. supervised the CD spectroscopy, and K.R. supervised the establishment of the production of recombinant PfKelch13 in Sf21 cells by R.S. and J.F.. R.S. and E.B. performed the growth curve analyses, protein purifications and DTNB assays. S.P. assisted with the cell culture, genotyping and western blot analysis. M.D. wrote the manuscript. All authors discussed the results and gave approval to the final version of the manuscript.

## Supplementary Information

**Supplementary Figure 1.**
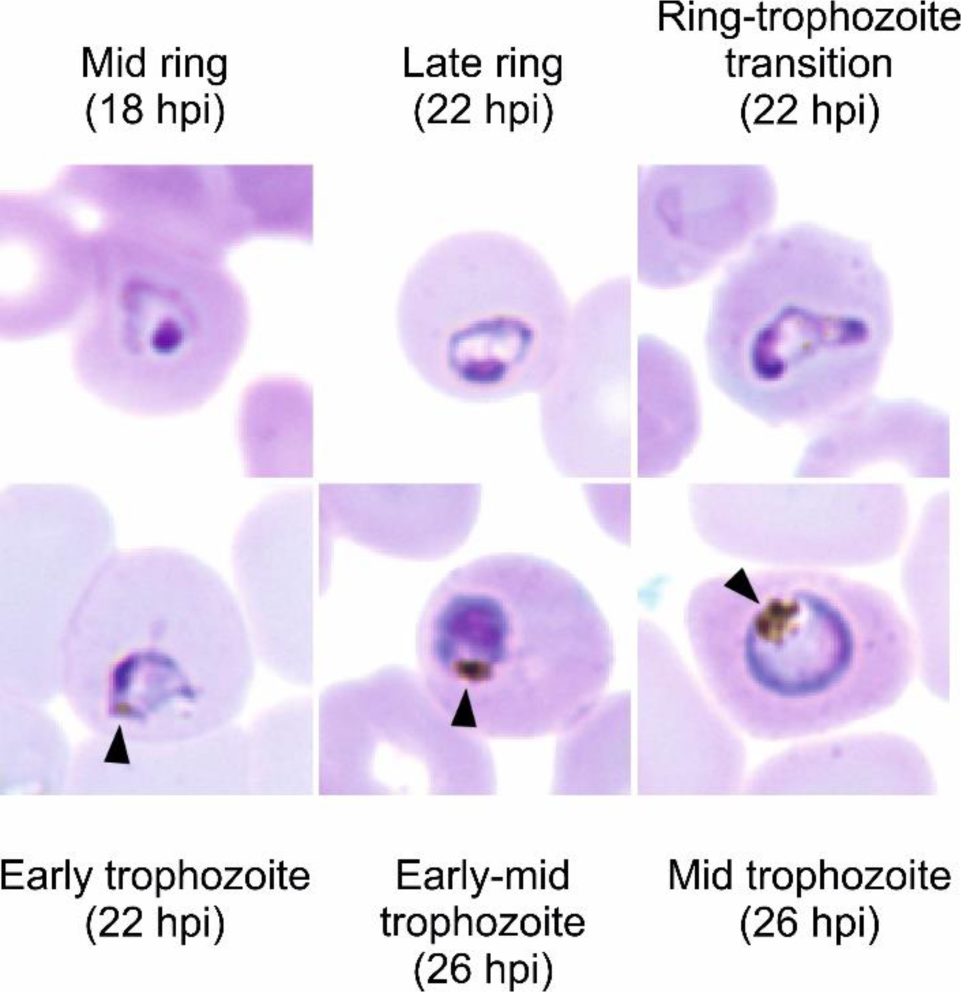
Representative images of stained blood-stage parasites. The hemozoin content of synchronized His_8_-*PFKELCH13-glmS* and His_8_-*PFKELCH13-M9* parasites was analyzed for air-dried and methanol-fixed thin blood smears that were stained with Hemacolor Rapid staining of blood smear solution. Images were acquired on a BZ-X800 microscope using the 60× oil objective. Time points when the cells were fixed are indicated. Visible hemozoin is labelled with an arrowhead.

**Supplementary Figure 2.**
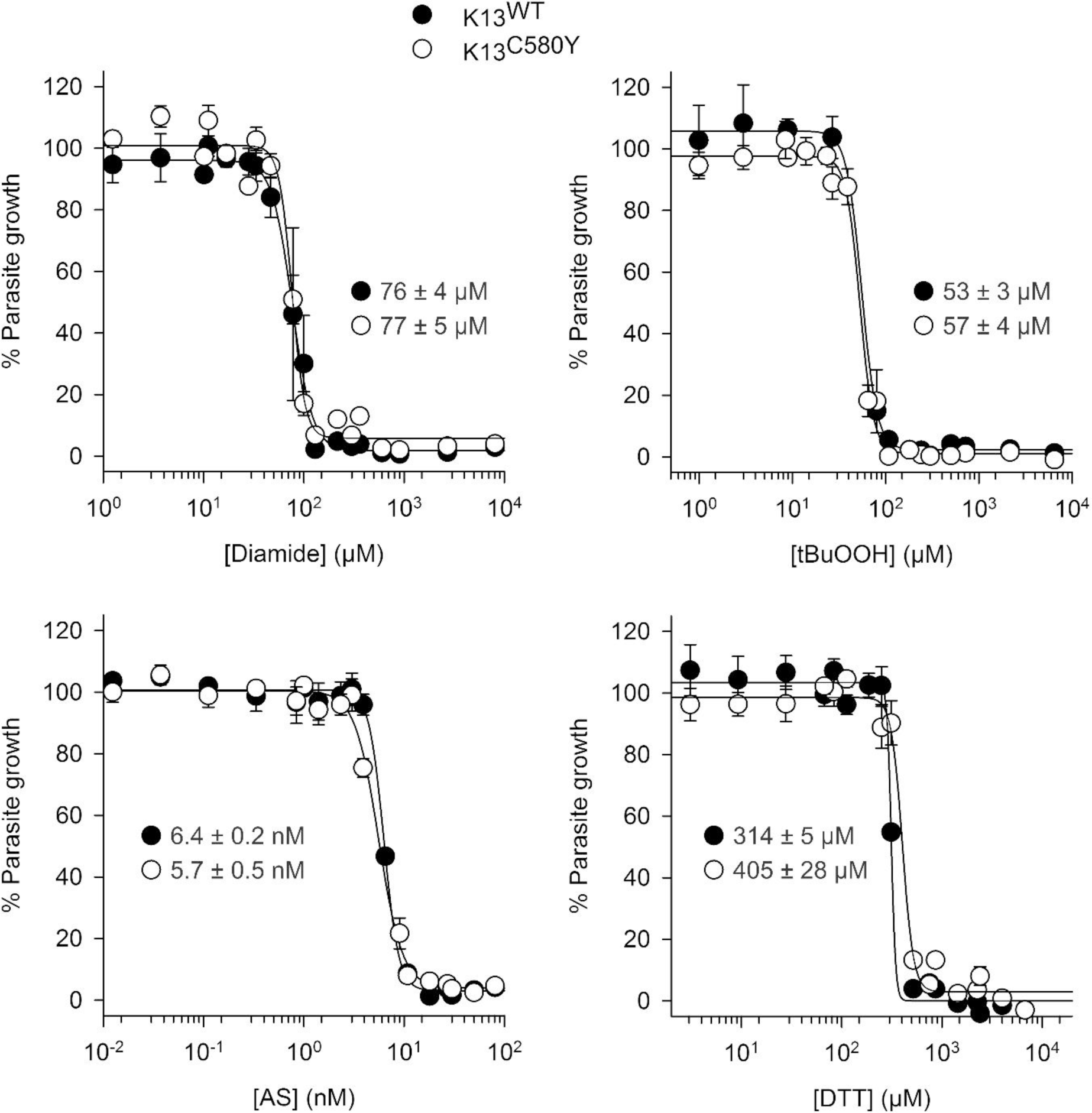
Growth inhibitory effects of redox agents on strain NF54K13^C580Y^. IC_50_ values were determined in 96-well plates using a SYBR green assay for bolus treatments of ring stage parasites with the indicated concentrations of the disulfide-inducer diamide, the oxidant tBuOOH, the endoperoxide artesunate (AS) and the reducing agent DTT. Each curve represents the mean ± standard deviation of three independent experiments with triplicate measurements. Data were fitted and analyzed in SigmaPlot13 revealing no significant differences between the indicated IC_50_ values for strains NF54K13^C580Y^ (K13^C580Y^) and wild-type NF54 (K13^WT^). Source data are provided as a Source Data file.

**Supplementary Figure 3.**
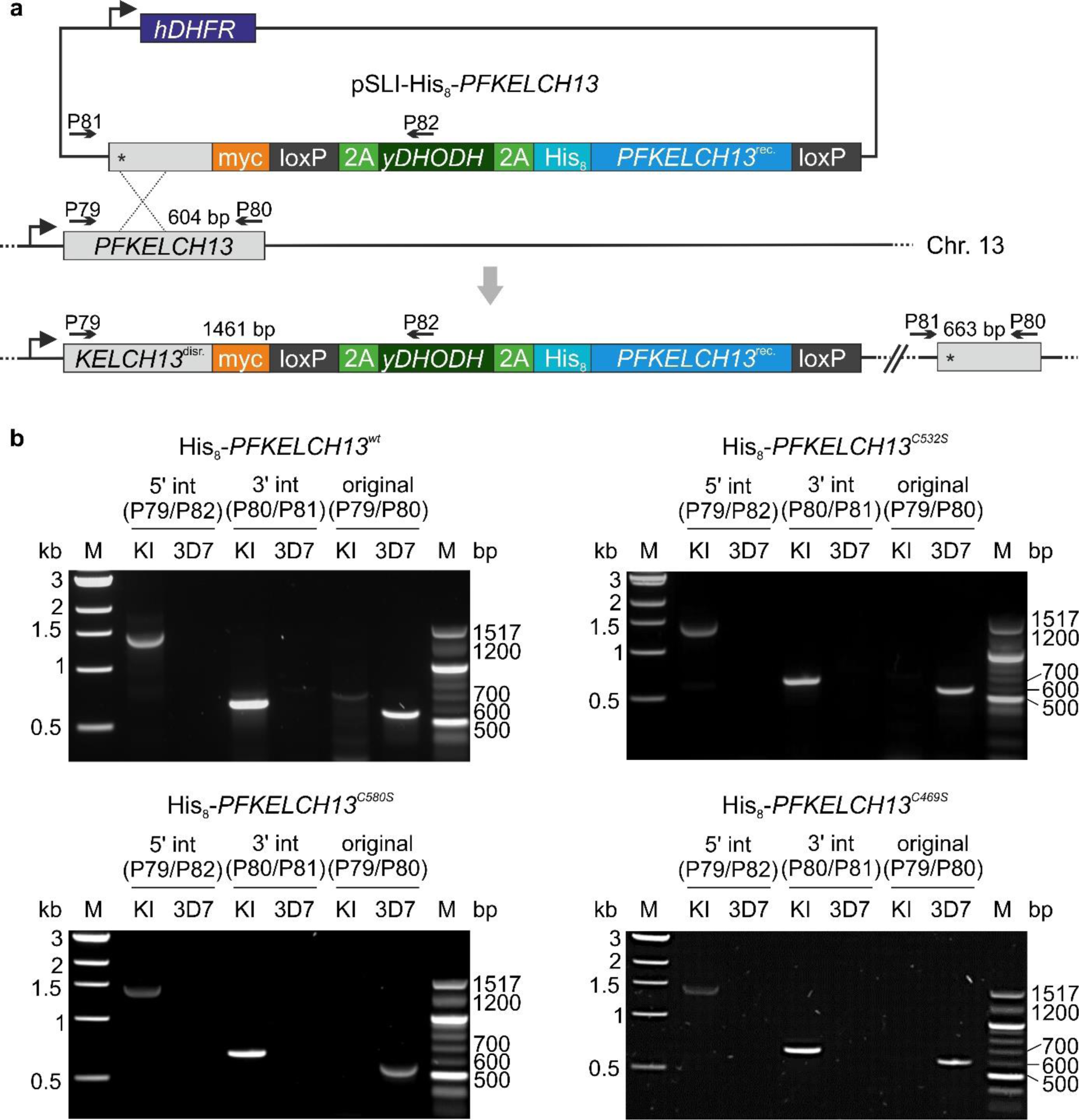
Generation and validation of *PFKELCH13* cysteine mutants. (**a**) Schematic overview of the SLI strategy. Recodonized *PFKELCH13* encoding His_8_-tagged wild-type protein or a cysteine mutant was fused with the selection marker *yDHODH* and integrated into *PFKELCH13* wild-type locus. Primer positions and expected product sizes for PCR analysis are highlighted. (**b**) PCR analyses using the indicated primer pairs from panel a confirmed the successful integration for wild-type strain His_8_-*PFKELCH13^wt^* and mutant strains His_8_-*PFKELCH13^C532S^*, His_8_-*PFKELCH13^C580S^*, and His_8_-*PFKELCH13^C469S^*. Genomic DNA from parental strain 3D7 served as a control.

**Supplementary Figure 4.**
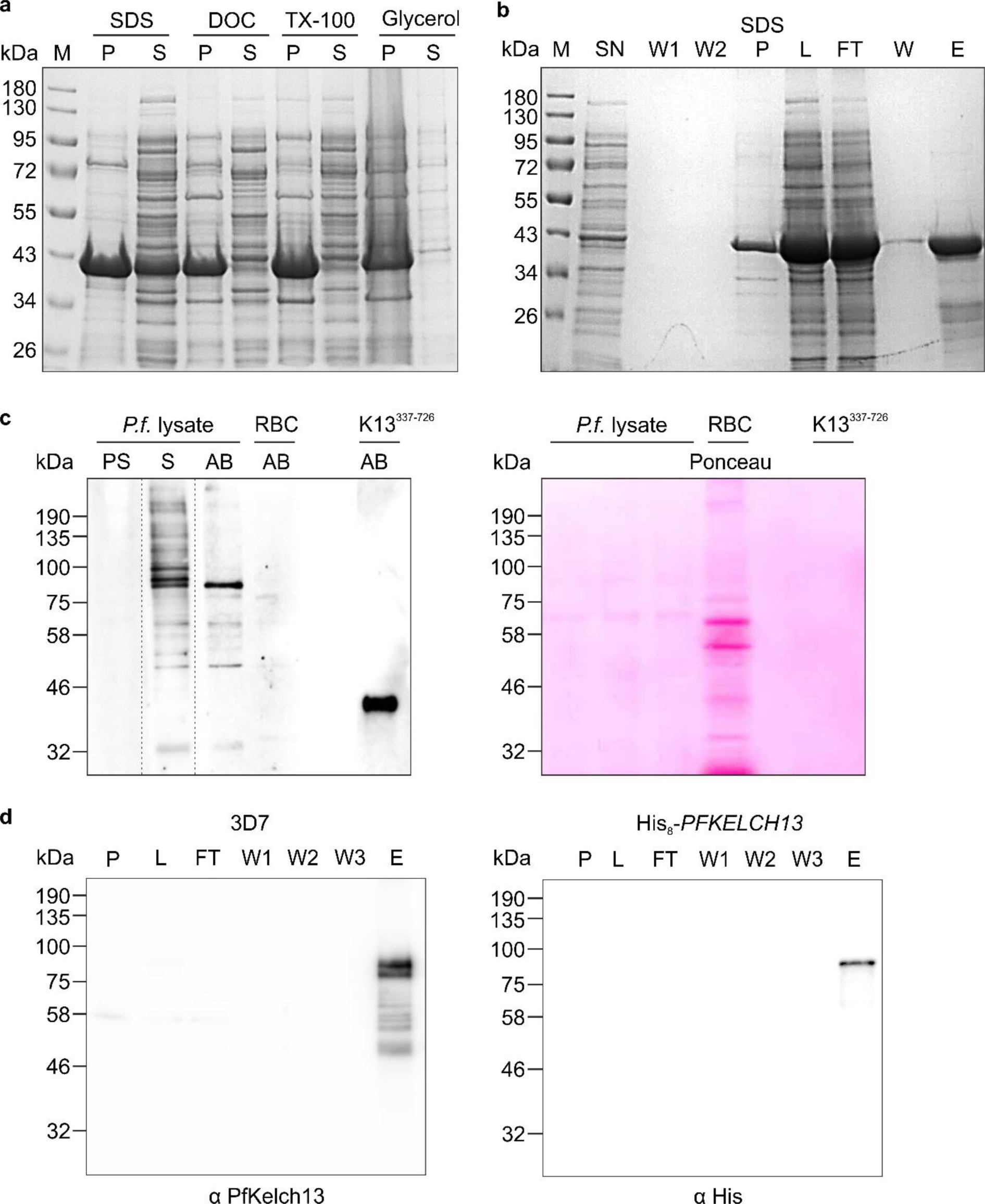
Generation of an antibody and pull-down of PfKelch13. (**a**) Solubilization screen for recombinant MAH_6_VGT-tagged PfKelch13^336-726^ from *E. coli*. PfKelch13^336-726^ was efficiently solubilized from washed inclusion bodies with solubilisation buffer containing 2% SDS, whereas 2% deoxycholic acid (DOC), 2% Triton X-100 (TX-100) or 50% glycerol did not show a solubilization effect. P: pellet, S: supernatant. (**b**) Purification of denatured recombinant PfKelch13^336-726^. Inclusion bodies containing PfKelch13^336-726^ were washed twice (W1, W2) and solubilized with 2% SDS. Excess SDS was precipitated on ice (SDS P). The supernatant (L) was loaded on a Ni-NTA agarose column. After washing (W), the protein was eluted with 200 mM imidazole (E) and used for immunization, yielding the serum and antibody in panel c. SN: supernatant from the *E. coli* lysate, FT: flow through. (**c**) Purification and validation of a rabbit PfKelch13 antibody by western blot analysis. Left: Serum (S) as well as purified antibody (AB) detected a protein with a molecular mass of ∼85 kDa in a *P. falciparum* lysate. Endogenous PfKelch13 has a calculated molecular mass of 83.7 kDa. Decoration with preserum (PS) served as a negative control. No signal was detected in uninfected red blood cells (RBC). Recombinant PfKelch13^337-726^ (K13^337-726^) with a calculated molecular mass of 45.7 kDa served as a positive control. Lanes contained either proteins from 3×10^7^ purified trophozoites, 3×10^7^ uninfected RBC or 10 ng PfKelch13^336-726^ as indicated. The dashed line indicates the cut membrane. Right: Ponceau S-staining of the membrane served as a loading control. (**d**) Purification of PfKelch13 from strain 3D7 (left) or His_8_-*PFKELCH13^wt^* (right) by immunoaffinity chromatography. The PfKelch13 antibody from panel c was coupled to CNBr-activated sepharose. Lysates with denatured proteins were prepared from 6x10^9^ trophozoites. P, pellet; L, 0.2% of the loaded lysate; FT, flow-through; W1-3, wash fractions 1-3; E, 6.7% of the eluate. The purified proteins were detected using either the PfKelch13 rabbit antibody or a commercial anti-His_6_ mouse antibody.

**Supplementary Figure 5.**
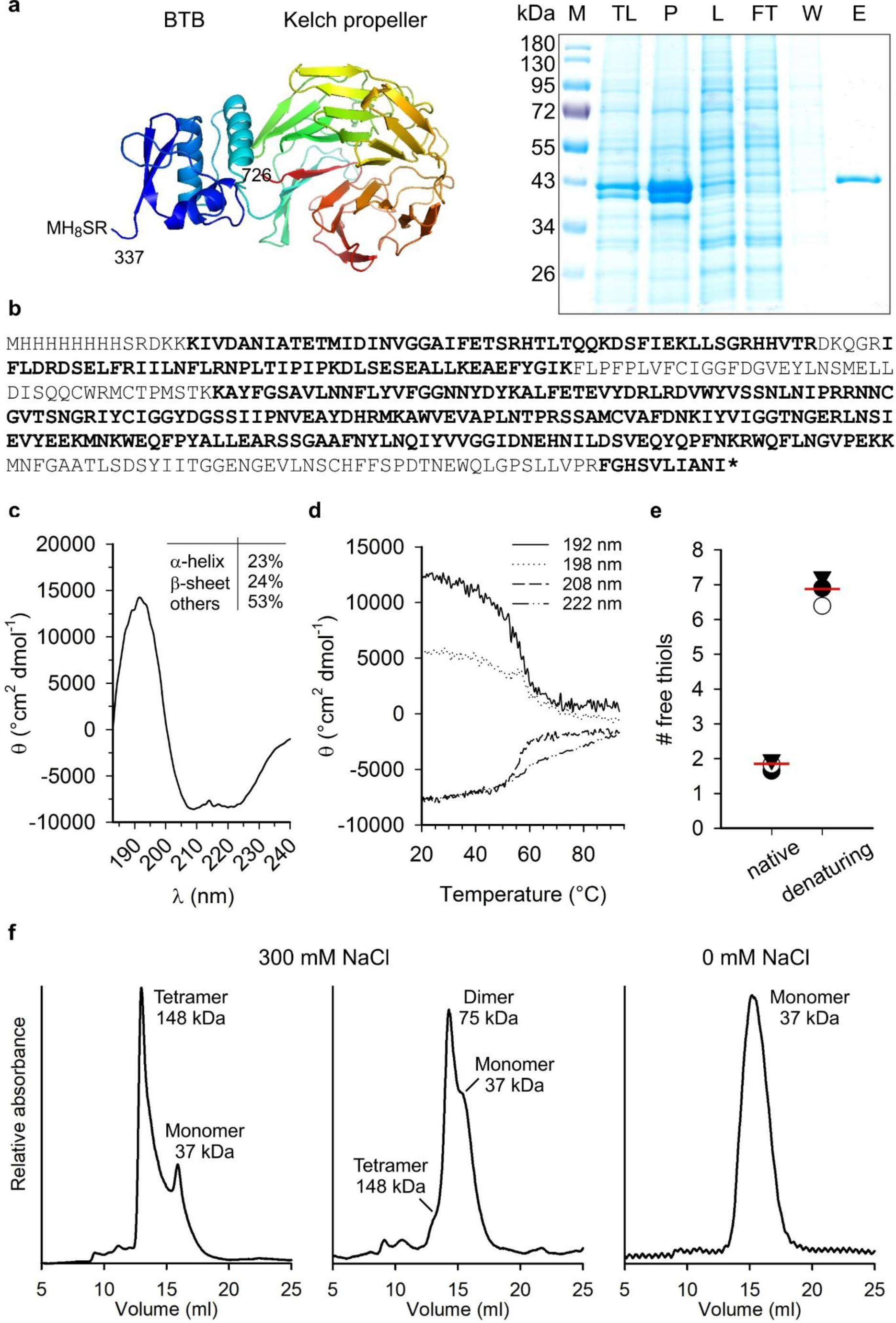
Purification and characterization of recombinant His_8_-PfKelch13. (**a**) Construct used for the production of MH_8_SR-tagged PfKelch13^337-726^ in Sf21 cells based on the released unpublished crystal structure (PDB entry 4YY8). Purification of recombinant PfKelch13^337-726^ by Ni-NTA affinity chromatography. Sf21 cells were lysed by three freeze-thaw cycles yielding the total cell lysate (TL). After removal of the insoluble material (P), the supernatant (L) was loaded on a Ni-NTA agarose column. The flow-through (FT) was discarded, the resin was washed (W) and the bound protein eluted (E) with 250 mM imidazole. (**b**) Sequence coverage of purified recombinant PfKelch13^337-726^ by mass spectrometry. Detected peptides are highlighted in bold. (**c**) CD spectrum of recombinant PfKelch13^337-726^ in 100 mM NaF, 50 mM Na_x_H_y_PO4, pH 7.4 at 20°C. Proportions of secondary structure elements were calculated using the K2D2 software. (**d**) Thermal stability of recombinant PfKelch13^337-726^ in the same buffer. (**e**) Detection of accessible cysteine residues of recombinant PfKelch13^337-726^ by DTNB in the absence and presence of 2% SDS. Data and mean from three independent measurements are shown. (**f**) Freshly purified recombinant PfKelch13^337-726^ was analyzed on a Superdex 200 Increase 10/300 GL column using 50 mM Na_x_H_y_PO_4_, pH 8.0 with (left) or without (right) 300 mM NaCl as running buffer at 10°C. The apparent molecular masses and interpreted oligomerization states are indicated.

**Supplementary Figure 6.**
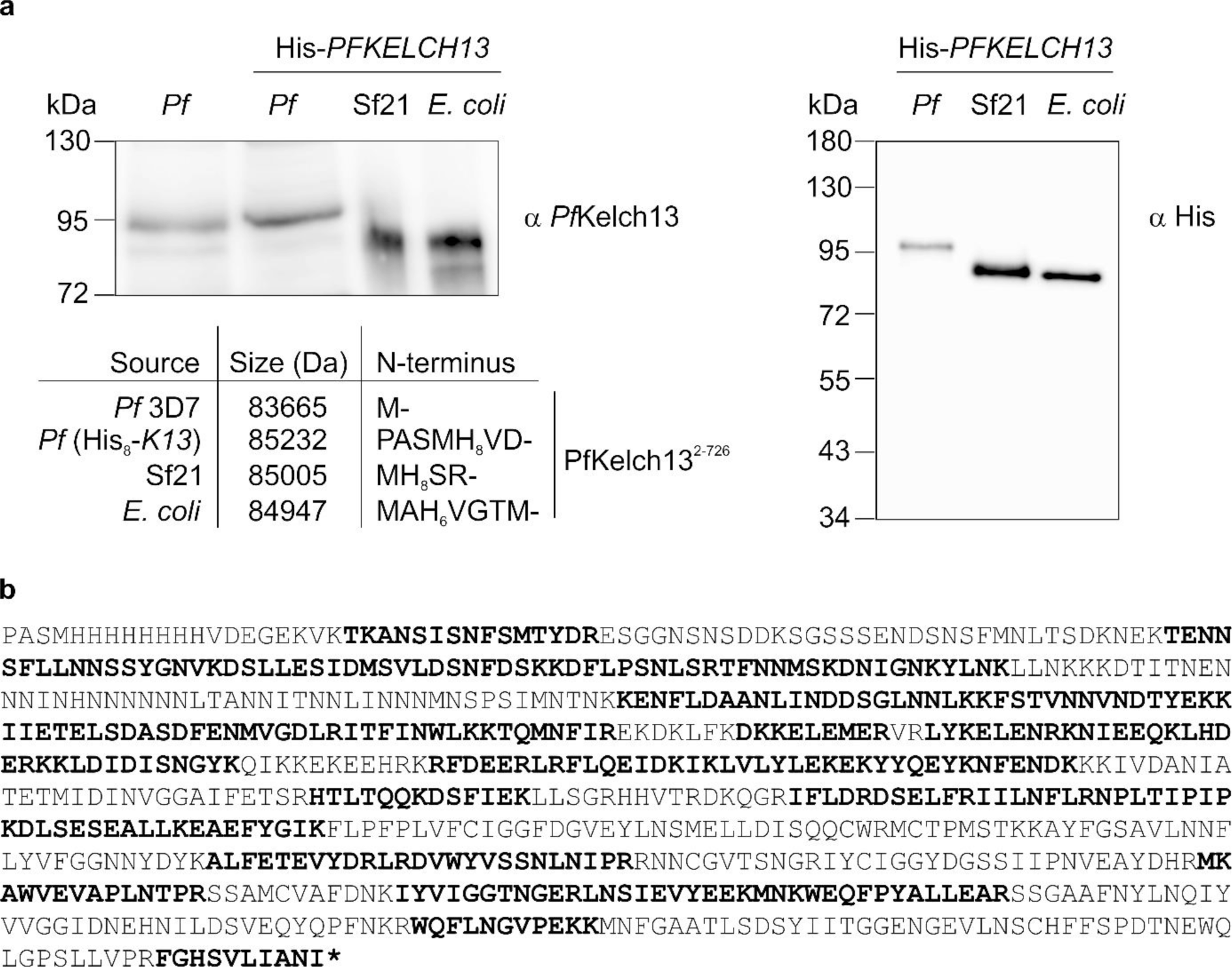
Potential post-translational modification of endogenous His_8_-PfKelch13. (**a**) N-terminally His-tagged PfKelch13 was produced in *P. falciparum* (*Pf*), *S. frugiperda* Sf21 cells (Sf21), and *E. coli*. Crude lysates were either probed with our PfKelch13 antibody from Fig. S3 (left) or an anti-His antibody (right). Crude extracts from strain 3D7 with wild-type PfKelch13 served as a control. Expected molecular masses and differences of the N-terminal tags before residues 2-726 are indicated. The PAS tripeptide at the N-terminus of His-tagged PfKelch13 from *P. falciparum* is derived from the cleaved 2A peptide. (**b**) Sequence coverage of His-tagged PfKelch13 that was purified by immunoaffinity chromatography from strain His_8_-*PFKELCH13^wt^* as shown in Fig. S3d. Detected peptides are highlighted in bold.

**Supplementary Figure 7.**
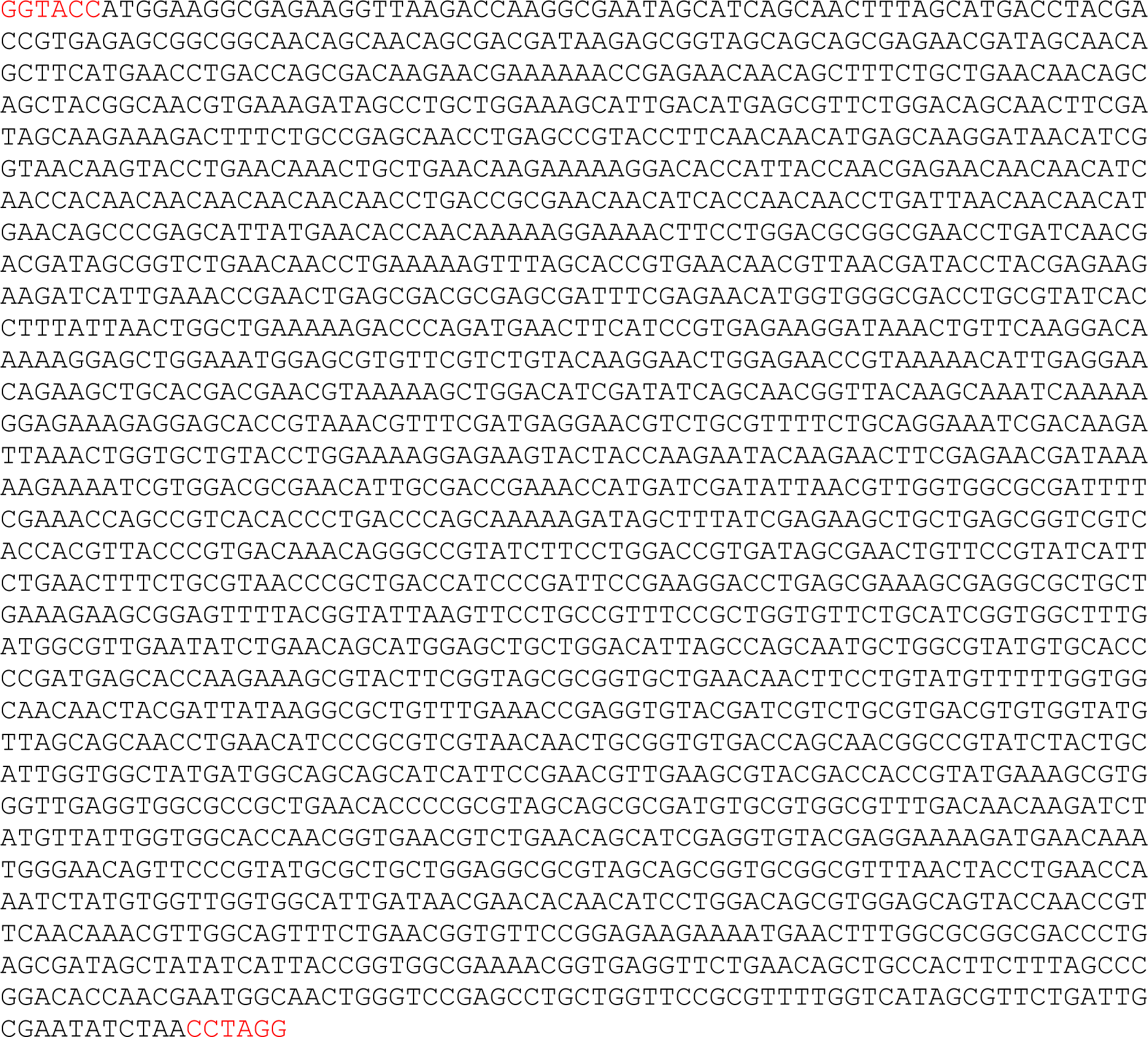
Sequence of *E. coli* codon-optimized *PFKELCH13*. The *Kpn*I and *Avr*II restriction sites of pET45b are highlighted in red.

**Supplementary Table S1.**
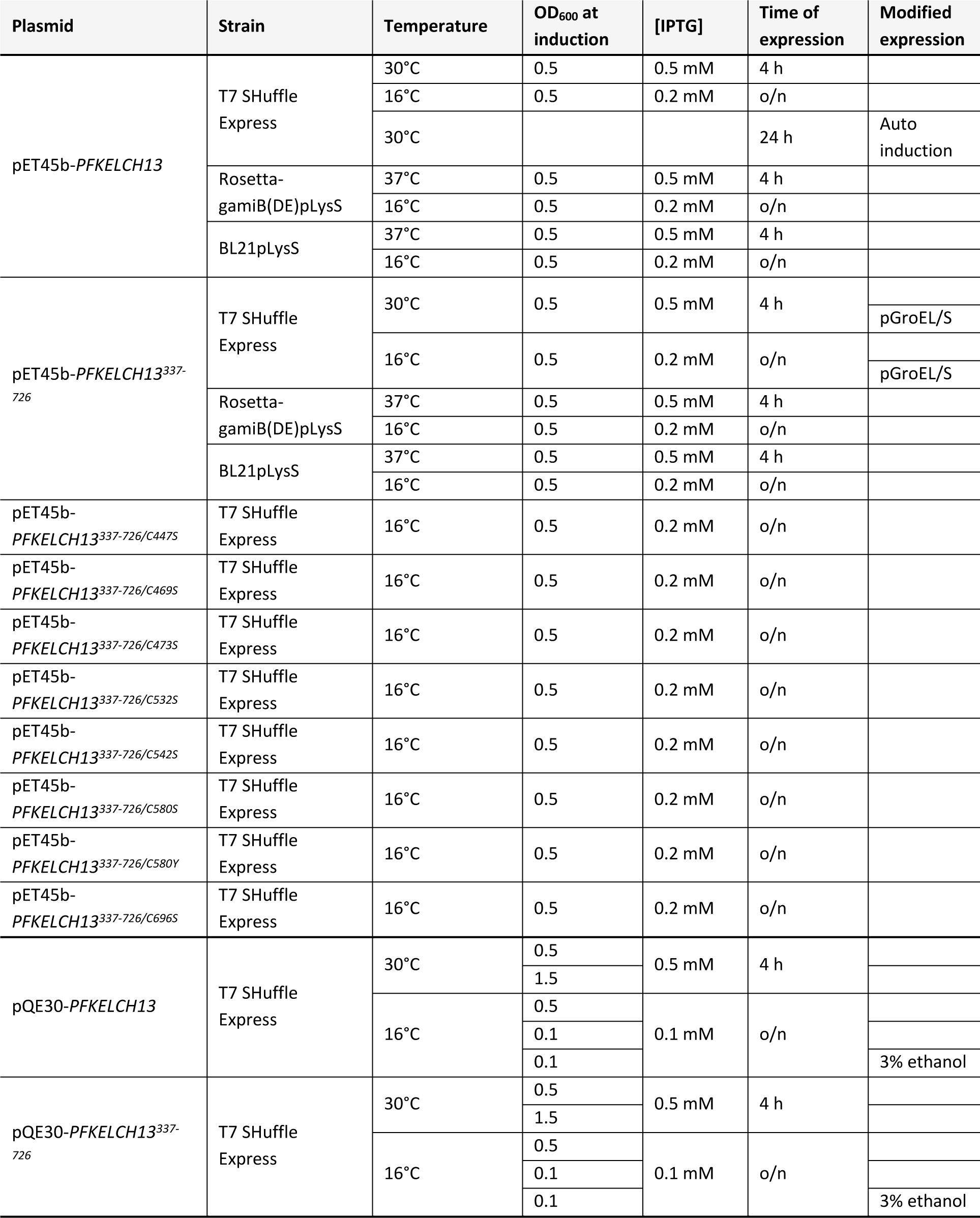
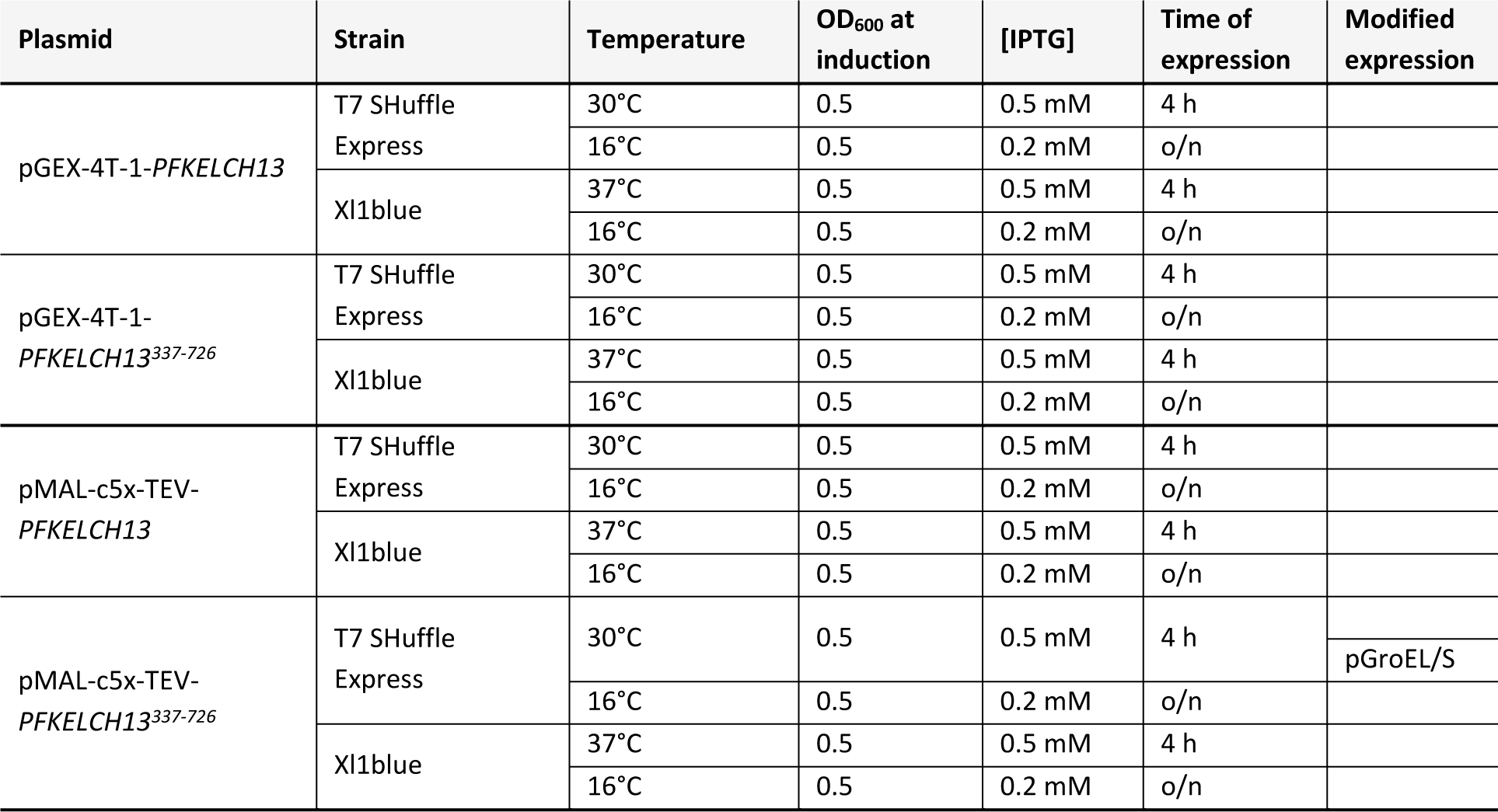
Tested conditions to produce recombinant PfKelch13 and PfKelch13^337-726^ in *E. coli*.

**Supplementary Table S2.**
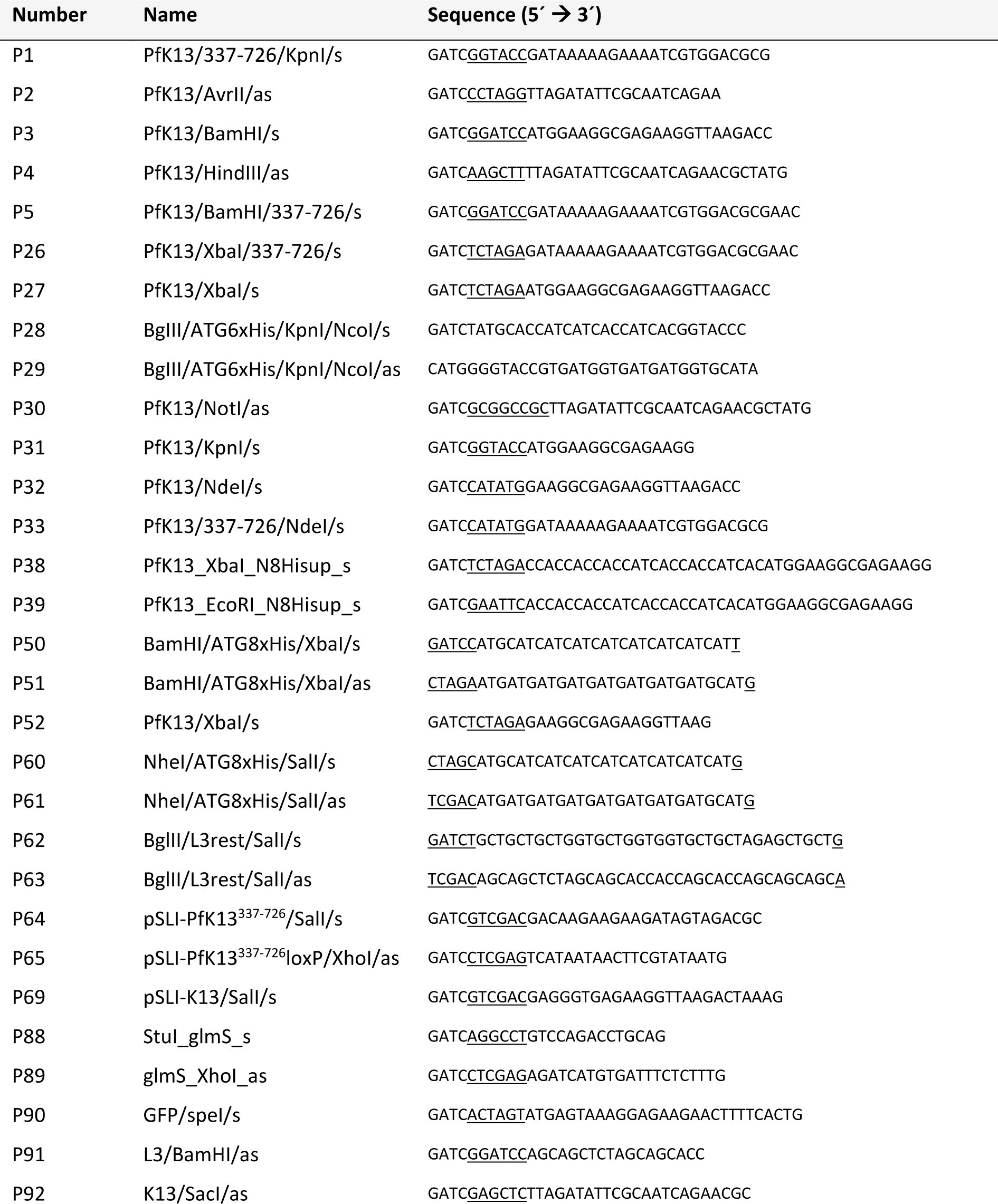
List of primers used for cloning of SLI and expression constructs. Restriction sites are underlined.

**Supplementary Table S3.**
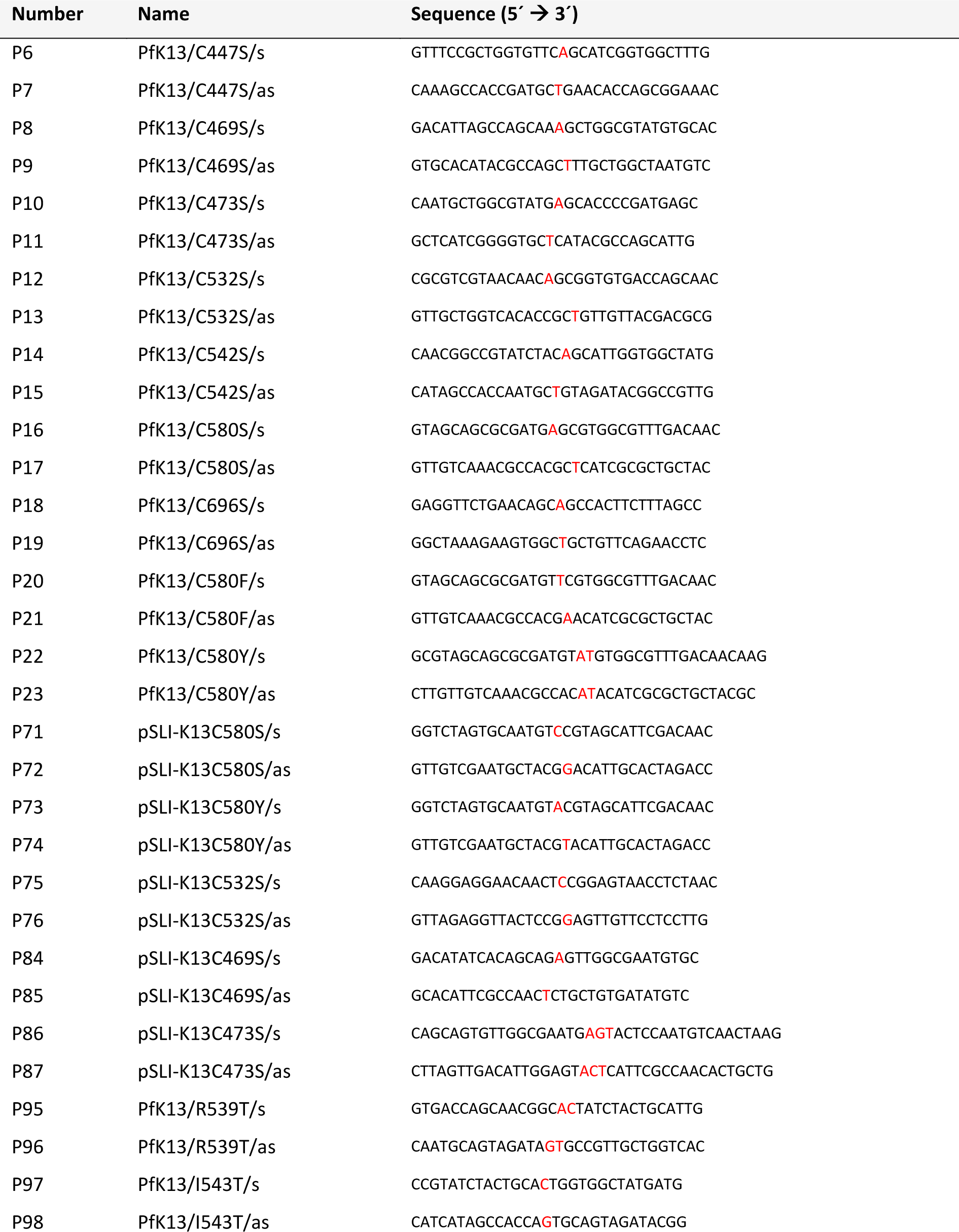
List of mutagenesis primers. Mutations are indicated in red.

**Supplementary Table S4.**
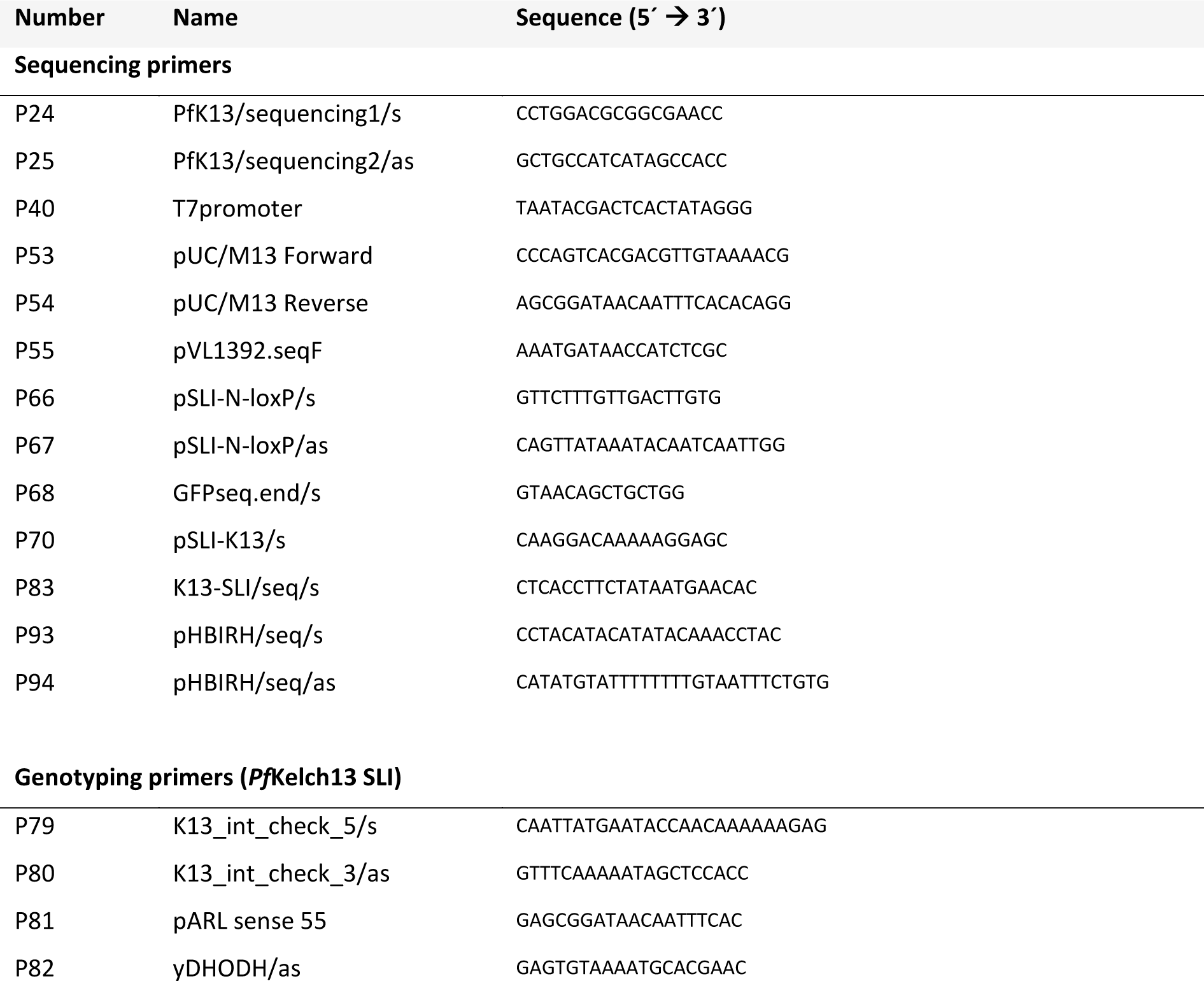
List of sequencing and genotyping primers.

